# Molecular organization of CA1 interneuron classes

**DOI:** 10.1101/034595

**Authors:** Kenneth D. Harris, Lorenza Magno, Linda Katona, Peter Lönnerberg, Ana B. Muñoz Manchado, Peter Somogyi, Nicoletta Kessaris, Sten Linnarsson, Jens Hjerling-Leffler

## Abstract

GABAergic interneurons are key regulators of hippocampal circuits, but our understanding of the diversity and classification of these cells remains controversial. Here we analyze the organization of interneurons in the CA1 area, using the combinatorial patterns of gene expression revealed by single-cell mRNA sequencing (scRNA-seq). This analysis reveals a 5-level hierarchy of cell classes. Most of the predicted classes correspond closely to known interneuron types, allowing us to predict a large number of novel molecular markers of these classes. In addition we identified a major new interneuron population localized at the border of *strata radiatum* and *lacunosum-moleculare* that we term “R2C2 cells” after their characteristic combinatorial expression of *Rgs12, Reln, Cxcl14*, and *Cpne5*. Several predictions of this classification scheme were verified using *in situ* hybridization and immunohistochemistry, providing further confidence in the gene expression patterns and novel classes predicted by the single cell data.

## Introduction

Interneurons play an essential role in governing the flow of information through neural circuits (Buzsaki and Chrobak, 1995, McBain and Fisahn, 2001, Markram et al., 2004, Fishell and Rudy, 2011, Isaacson and Scanziani, 2011, Somogyi et al., 2014). Cortical circuits contain a multitude of GABAergic interneuron classes, characterized by diverse axonal and dendritic structure, intrinsic electrophysiological properties, connectivity, *in vivo* firing characteristics, as well as developmental history and gene expression. The organization of these classes is partially, but not completely preserved between different cortical structures such as hippocampus, amygdala, and isocortex (Klausberger and Somogyi, 2008, Petilla Interneuron Nomenclature et al., 2008, Rudy et al., 2010, Spampanato et al., 2011, Bartolini et al., 2013, Defelipe et al., Pfeffer et al., 2013, Kepecs and Fishell, 2014, Somogyi et al., 2014, Jiang et al., 2015). Yet while the diversity of cortical interneurons has been studied in great detail, their classification remains controversial (Defelipe et al., 2013). Understanding the diversity of interneurons thus remains a challenging goal for neuroscience.

Interneurons have been studied in great depth in area CA1 of the hippocampus, where they have been classified into at least 21 distinct types characterized by neurochemical, connectional, and firing patterns (Freund and Buzsaki, 1996, Klausberger and Somogyi, 2008, Somogyi, 2010, Bezaire and Soltesz, 2013). This work has revealed the great diversity and intricate organizing principles that can exist for interneurons of a single brain region. Moreover, the diversity revealed so far is likely to be incomplete: there are almost certainly further subdivisions of the classes established so far, as well as potential “dark matter” classes that have to date escaped molecular identification, as the molecular markers that would identify them are unknown. For example, neurons at the border of stratum radiatum and lacunosum-moleculare (R-LM cells) form a diverse population which play an important role in integrating multiple inputs and outputs of the CA1 region (Vida et al., 1998, Miyashita and Rockland, 2007, Melzer et al., 2012, Kitamura et al., 2014), but whose molecular organization is to-date poorly understood.

One of the principal characteristics used to define interneuron classes is their molecular fingerprint: the set of genes they transcribe to mRNA and translate to protein. The molecular code identifying interneuron classes is likely to be complex and combinatorial, and it appears unlikely that there exists a one-to-one correspondence between interneuron classes and individual molecular identifiers. In CA1, for example, the calcium binding protein *Pvalb* is expressed without the neuropeptide *Sst* in fast-spiking basket cells, *Pvalb* and *Sst* are strongly expressed together in bistratified cells, while other interneuron classes express *Sst* strongly but *Pvalb* only at lower levels (Katona et al., 2014). Similarly complex combinatorial relationships hold with many other markers (Somogyi, 2010, Bezaire and Soltesz, 2013, Wheeler et al., 2015). Traditional methods of molecular histology – which can identify at most a handful of markers simultaneously – are thus unlikely to reveal the full complexity of interneuronal classes. Indeed, some authors have suggested that interneuronal diversity is so extreme that a continuum of properties might be a more accurate description than discrete classes (Parra et al., 1998, Markram et al., 2004).

The new technique of single-cell RNA sequencing (scRNA-seq), which quantifies the mRNA expression levels of all genes in each studied cell, provides an unprecedented opportunity to identify cell types by their combinatorial expression of molecular markers (Jaitin et al., 2014, Pollen et al., 2014, Treutlein et al., 2014, Grun et al., 2015, Macosko et al., 2015, Usoskin et al., 2015, Zeisel et al., 2015). To fully harness the data that comes from this method, however, requires sophisticated computational analysis. A cell’s mRNA expression profile can be considered as a vector in an approximately 20,000 dimensional space. Classification of cells is a form of “cluster analysis”: identifying sets of vectors which are similar according to an appropriate criterion. High-dimensional cluster analysis is a notoriously difficult problem (Bouveyron and Brunet-Saumard, 2014), and while multiple algorithms have been proposed, algorithms must typically be tailored specifically to individual problems. Cell typing from scRNA-seq is made more challenging by the fact that not all variations in mRNA expression reflect differences in cell type: as well as potential methodological artifacts, gene expression levels change dynamically (Kaern et al., 2005), and are modulated by conditions such as sleep, activity, and learning (Cirelli et al., 2004, Donato et al., 2013, Dehorter et al., 2015). Thus, to be certain that scRNA-seq cell typing is producing accurate results, it is essential to calibrate this method in a system where cell classes have already been extensively studied. Indeed, it is only after verifying that a cell-typing algorithm correctly identifies cell classes previously defined by traditional methods that one can be confident in the prediction of novel classes found by the same algorithm.

Here, we use scRNA-seq data to characterize the molecular organization of CA1 interneurons, revealing a 5-level hierarchy of cell types. This analysis reveals a major and previously uncharacterized molecular class of interneuron, characterized by expression of novel marker genes including *Rgs12, Reln, Cxcl14*, and *Cpne5*. The spatial expression patterns of these genes allow us to locate the class at the border of *strata radiatum* and *lacunosum-moleculare*, a region containing incompletely-characterized but functionally important interneurons (Vida et al., 1998, Miyashita and Rockland, 2007, Melzer et al., 2012, Kitamura et al., 2014). Other branches of the hierarchy show expression patterns closely matching those of known interneuron classes, providing confidence in the novel classes identified by the algorithm, and revealing a rich set of predictions for novel molecular markers of these classes.

## Results

We analyzed a database of 126 cells dissociated from mouse area CA1 (CD-1 strain, ages P21-P31, both sexes), and subjected to scRNA-seq and identified as interneurons using previously described methods (Zeisel et al., 2015). The starting point of the current study was thus a 126 by 19,972 matrix of integers, giving the number of detected RNA molecules for each gene in each interneuron.

Before describing our algorithm and results, it is important to consider two technical artifacts that can arise with scRNA-seq. First, the fractions of each cell type that are sequenced need not correspond to their abundance in native tissue, due to differing survival probabilities of different cell classes during tissue dissociation. For example, only ~5% of the cells in the current database strongly express *Pvalb* whereas immunohistochemical analysis suggests this fraction should be closer to ~20% (Jinno and Kosaka, 2006). Second, a minority of sequenced cells show contamination by RNAs of another cell class, due to cell-cell adhesion during the dissociation phase. For example, in a separate database of CA1 pyramidal cells (Zeisel et al., 2015), approximately 5% were positive for multiple oligodendrocyte markers, even though these genes are not truly expressed by pyramidal cells.

Manual examination of RNA expression levels revealed that different types of gene showed different patterns of expression (Figure 1). Consider *Sst*, a selective marker of a subset of interneurons. The histogram of expression levels of this gene across cells showed an extremely skewed distribution (Figure 1, top left panel), with a majority of cells expressing zero copies of the gene, but a smaller population of cells showing high levels. Other selective markers (e.g. *Cck, Cnr1, Pvalb;* further diagonal panels in Figure 1) showed similarly skewed expression histograms. By contrast, genes that are not markers of specific interneuron subtypes (such as *Actb*, universally expressed in all cells, and *Gad1*, expressed in all interneurons; bottom-right two diagonal panels in Figure 1) showed distributions without a prominent peak at zero.

**Figure 1.**
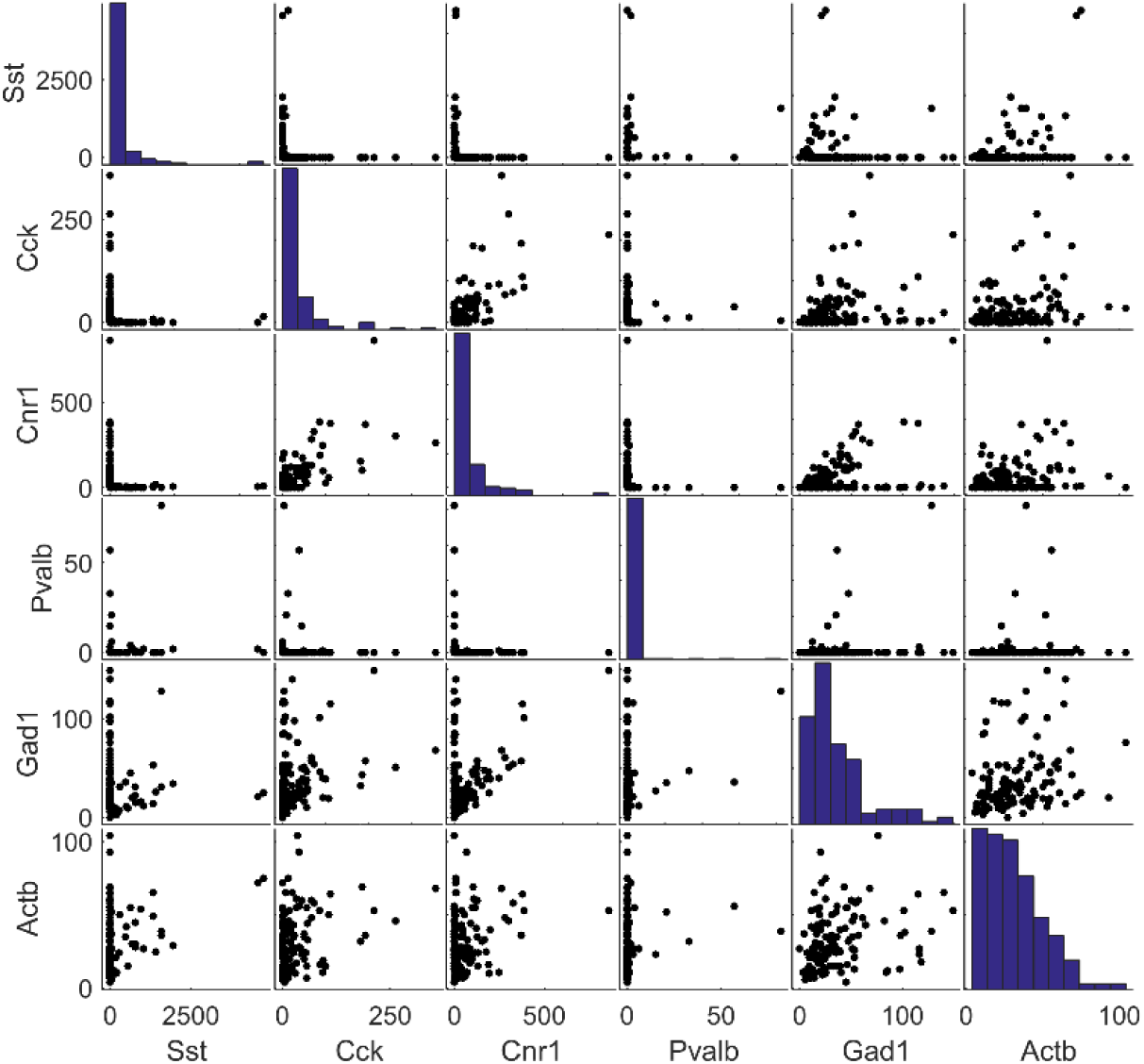
Scatter plot matrix showing expression levels of four classical interneuron marker genes (*Sst, Cck, Cnr1, Pvalb*), and two genes universally expressed in all interneurons (*Gad1, Actb*). Panels along the diagonal show histograms of expression for each gene; each panel off the diagonal shows scatterplots of expression for a particular gene pair, with each point representing a single cell.

Some pairs of marker genes showed mutually exclusive expression. For example, previous work suggests that the neuropeptides *Sst* and *Cck* are expressed in distinct subpopulations of CA1 interneurons (Somogyi, 2010, Bezaire and Soltesz, 2013). Our scRNA-seq data was consistent with this result: while many cells expressed one of these two genes at high levels, and many expressed neither, no cell strongly expressed both together. A scatter plot of expression of these two genes thus resembled an uppercase letter “L” (Figure 1). While expression of *Sst* and *Cck* was almost completely mutually exclusive, other pairs of genes showed weaker forms of exclusivity. For example, while *Pvalb* expression was generally weak in *Sst* positive cells, it was not always zero, consistent with prior immunohistochemistry (Katona et al., 2014); and while *Cck* expression was generally weak in *Pvalb* neurons, it was not absent, consistent with its previous detection in RT-PCR experiments (Tricoire et al., 2011).

Some pairs of marker genes showed positive correlations, suggesting that they are expressed in the same interneuronal populations. For example, previous work has suggested that the cannabinoid receptor gene *Cnr1* is expressed in most Cck-positive interneurons (Katona et al., 1999). Our scRNA-seq data was consistent with this result, showing that expression levels of *Cck* and *Cnr1* were positively correlated (Figure 1).

Universally-expressed genes showed a different pattern of expression and correlation compared to marker genes. For example, *Gad1* and *Actb* showed wide ranges of expression but not strong peaks at zero; furthermore, while weak positive correlations were often seen between these genes, mutual exclusivity was not observed. Scatter plots of universally-expressed genes against marker genes typically showed a “chevron-shaped” distribution, with correlations seen within the population of cells that express the marker gene (e.g. *Gad1* vs. *Pvalb)*.

We conclude that mRNA expression levels are diverse between cells and can be correlated, but that this diversity cannot cause simultaneous expression of two mutually-exclusive marker genes. We therefore designed an algorithm to search for groups of marker genes with mutually-exclusive expression patterns, and use them to classify cells in a manner that is robust to variability in their absolute expression levels.

## Splitting cells into branches by finding teams of marker genes

The algorithm we designed works by searching for mutually exclusive “teams” of genes, such that each cell expresses genes from one team or the other, but not from both. Specifically, a gene team is defined by a *N_genes_*-dimensional non-negative weight vector **w**. The mRNA expression levels of a cell are summarized in a *N_genes_*-dimensional vector **x**, and the cell is assigned a “team score” given by the scalar product **w** · **x**. The algorithm searches for a pair of weight vectors that optimize an objective function which is large when each cell has one but only one team score larger than zero, the weight vectors are sparse, and the cells are divided into branches of roughly equal size (full details provided in Experimental Procedures).

We applied the algorithm in a recursive manner. At the top level of the resulting classification hierarchy, the algorithm split the CA1 interneurons into two branches containing 57 and 69 cells. Each cell had one, but only one team score greater than zero, as required to optimize the objective function (Figure 2a). The weight vectors identified had 6 and 10 non-zero entries, which defined the corresponding top-level gene teams (Figures 2b, 2c). The split found by the algorithm was statistically significant: when the same algorithm was applied to an ensemble of randomized gene expression matrices (see Experimental Procedures), mutually exclusive gene teams were not found, resulting in substantially lower objective function values (Figure 2d). Examination of a scatterplot matrix of gene expression values (Figure 3) confirmed that the gene teams were mutually exclusive: cells expressing at least one gene from team 1 did not express any genes from team 2, and *vice versa*.

**Figure 2.**
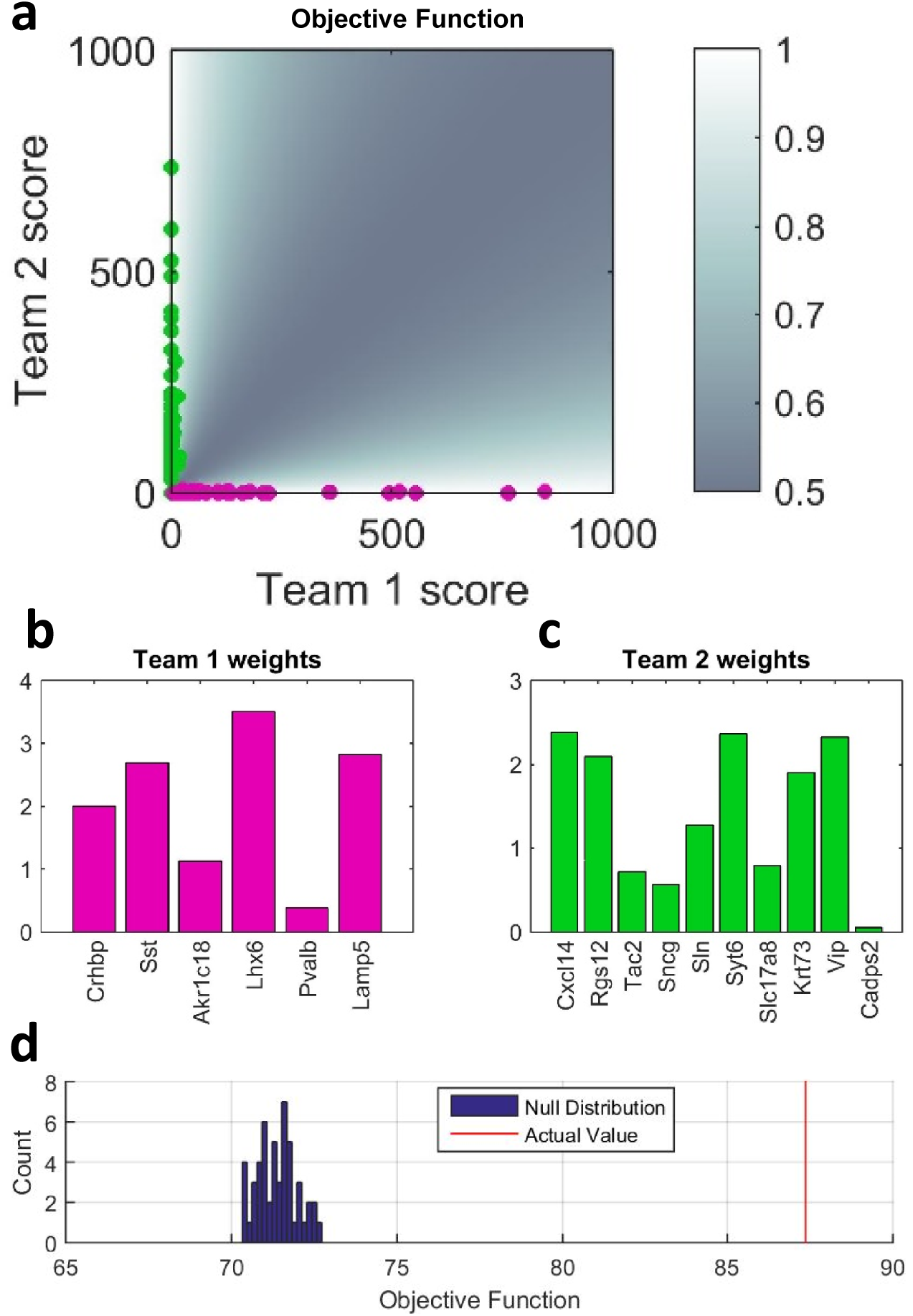
The GeneTeam algorithm splits a set of cells into two branches by finding two teams of genes, such that each cell expresses genes from one but only one team. (**a**), Scatter plot showing the team scores (i.e. weighted-sum expression levels) underlying the top-level division of CA1 interneurons. Magenta and green points represent cells classified into branches 1 and 2. Superimposed pseudocolor map shows the objective function whose sum is maximized by the algorithm; lighter colors indicate the preferred region in which one and only one team is expressed strongly. (**b, c**), Bar-chart representation of the weights for each gene in teams 1 and 2. (**d**), Statistical significance was assessed by comparing the optimal objective function to a null distribution computed on an ensemble of randomized gene expression matrices. The actual value of the objective function is far outside the null distribution, indicating that the split was statistically significant.

**Figure 3.**
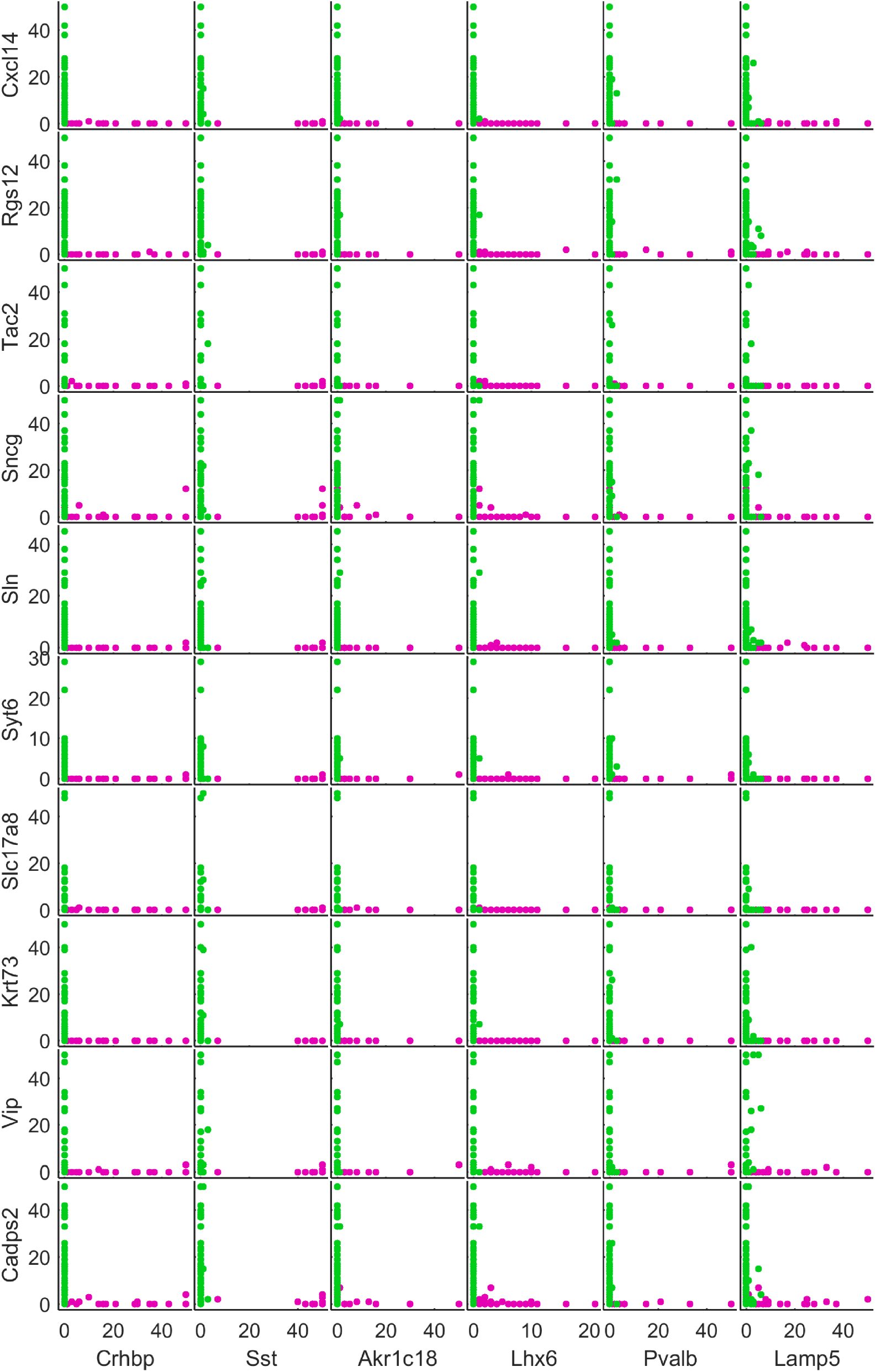
Scatterplot matrix showing expression of each pair of genes from the two top-level teams. Magenta and green dots indicate cells assigned to branches 1 and 2. Note that a cell assigned to branch 1 cannot strongly express any genes from team 2, and *vice versa*. Expression levels for each gene have been clipped to a maximum of 50 molecules/cell to aid visibility.

Some of the genes defining the top-level split were familiar interneuron markers (such as *Sst, Lhx6, Pvalb, Slc17a8* and *Vip*), but others were novel. Although the algorithm assigns cells into branches based solely on expression of gene team members, there are many other genes whose expression differs between branches. Examination of a scatterplot matrix for a subset of these genes (Supplementary Figure 1) showed that their expression differed between branches, but without the strict mutual exclusivity characteristic of gene team members. For example, while the serotonin receptor *Htr3a* was primarily associated with branch 2, a small but noticeable subset of branch 1 neurons expressed *Htr3a* at moderate levels. We conclude that CA1 interneurons can be divided into two top-level groups characterized by mutually-exclusive gene teams, and that additional genes whose expression differs between branches might provide information to further subdivide these branches at deeper levels.

Consideration of the specific genes whose expression differed between the two top-level branches suggested a biological interpretation. Cortical interneurons are developmentally derived from two primary sources, the medial and caudal ganglionic eminences (MGE and CGE). In isocortex, most interneuron types comprise cells with a unique origin in one of these two areas, and the developmental origin of a cell can be identified by its expression of *Htr3a*. In hippocampus however, some well-defined interneuron classes (such as neurogliaform and O-LM cells) contain subpopulations positive and negative for *Htr3a*, which have been suggested to reflect cells derived from CGE and MGE, respectively (Tricoire et al., 2010, Chittajallu et al., 2013). The 1^st^ level gene teams found by the algorithm generally matched markers of developmental origin: genes frequently expressed by cells in the first top-level branch (including *Sst, Pvalb, Lhx6, Satb1)* are most often associated with MGE-derived cells, whereas genes frequently expressed by cells in the second top-level branch (including *Vip, Cck, Slc17a8, Cnr1, Htr3a)* are more often associated with CGE-derived classes. We refer to these branches as “MGE-like” and “CGE-like”, to indicate that their adult gene expression patterns largely match those expected from MGE-derived and CGE-derived interneurons. Nevertheless, we note that each branch may also contain cells of the opposite developmental origin that have adopted this expression pattern at some point before adulthood. In addition to the classical markers identifying these teams, we found a number of novel markers including *Hapln1, Nxph1, Adcy1, Grin3a, Sparcl1, Rgs17, Ncald* expressed in subsets of the MGE-like branch, and *Npas1, Fxyd6, Trp53i11, Cadps2, Rgs12, Cxcl14, Sncg, Cplx2* expressed in subsets of the CGE-like branch (Supplementary figure 1).

The GeneTeam algorithm was applied recursively, resulting in a 5-level hierarchical classification tree (Figure 4). We next set out to compare the classes and expression patterns predicted by the GeneTeam algorithm to those previously established by immunohistochemistry, reasoning that if some of the algorithmically-defined classes showed close similarity to categories defined with established methods, this would build confidence in novel classes suggested by the algorithm. Although the results of this classification differed in detail to those produced by a previously described BackSPIN algorithm (Zeisel et al., 2015) (Supplementary Figure 2), we found a remarkably detailed level of correspondence to previous immunohistochemical and RT-PCR studies, allowing us to identify a large number of branches of the classification tree (Figure 4) and to predict the gene expression patterns of each class (shown for a selection of genes in Figure 5). While most branches of the classification tree closely matched previously identified cell classes (Figure 4, blue boxes), one branch could not be readily identified, and we assumed this to represent novel cell classes (Figure 5, green boxes). The reasoning behind these associations is presented next.

**Figure 4.**
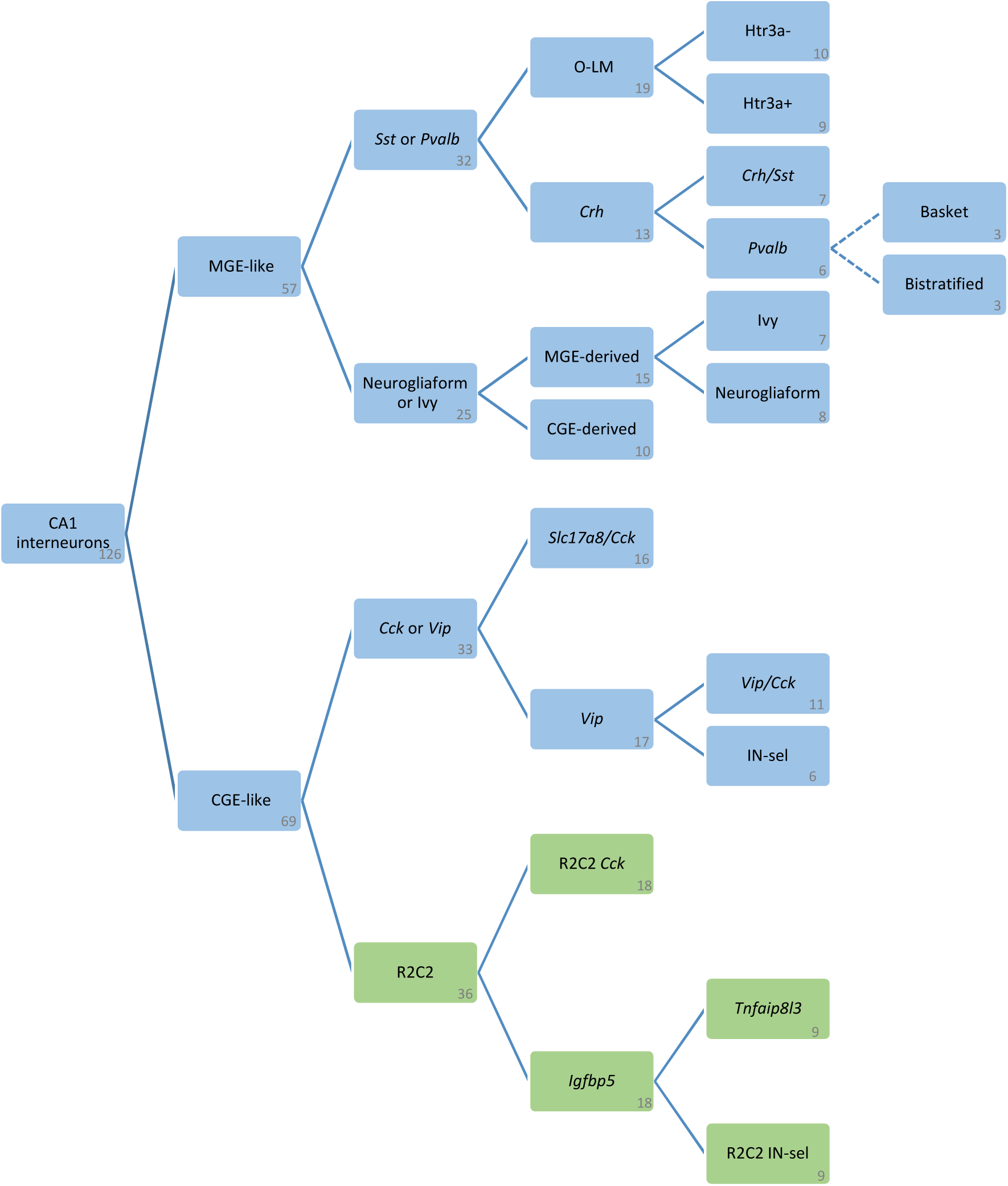
Classification tree produced by recursive application of the GeneTeam algorithm. Text labels indicate putative class of each branch; the number of cells in each branch is shown in the bottom right corner. Green boxes mark the novel R2C2 branch. The major statistically significant branch splits (*p* <10^−14^) are shown as solid lines; an additional split that did not reach significance, but was still judged as reflecting separate classes due to a correspondence with previous results, is shown by dashed lines.

**Figure 5.**
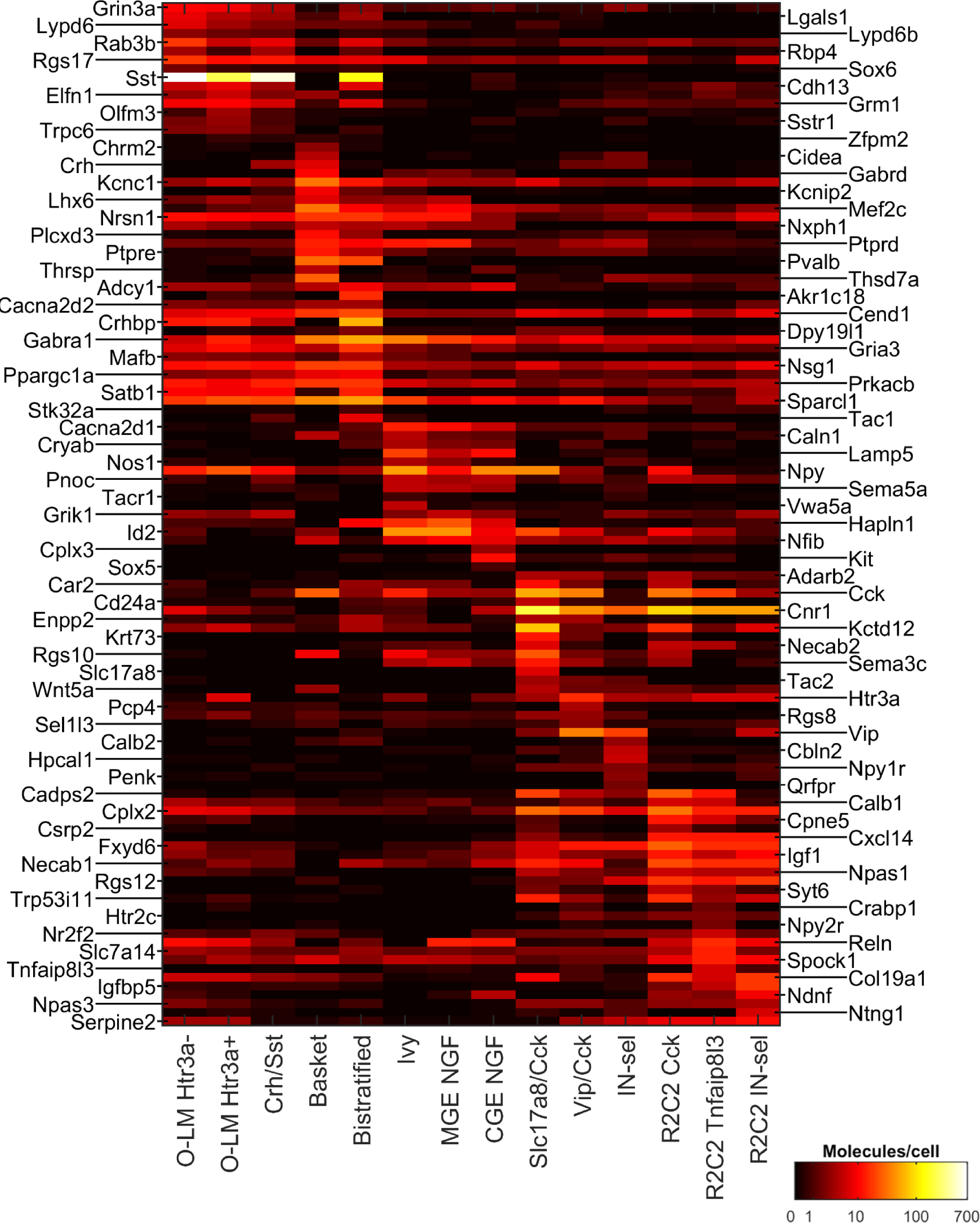
Heatmap showing mean expression level of selected genes in each putative cell class.

## *Sst* and *Pvalb* expressing interneurons

To aid the identification of each branch we developed a compact representation, in which the relative expression of selected genes for each branch is shown in pseudocolor format (Supplementary Figure 3). The MGE-like branch split into two 2^nd^-level branches, the first of these which was characterized by expression of *Sst* and *Pvalb*. We started by examining this branch in detail.

At the 3^rd^ level, the *Sst/Pvalb* branch contained a first branch that matched the expression profile of O-LM neurons, an Sst-positive type found in stratum oriens (Supplementary Figure 4). Indeed, this branch expressed *Reln* strongly and *Pvalb* at most weakly, and also expressed other O-LM markers including *Grm1* (Ferraguti et al., 2004), *Elfn1* (Sylwestrak and Ghosh, 2012), *Lhx6* (Liodis et al., 2007), *Satb1* (Close et al., 2012, Chittajallu et al., 2013), and *Lgals1* (Kajitani et al., 2014). In addition, this branch expressed several other genes including *Lypd6, Lypd6b, Crhbp, Grin3a*, and *Rab3b*. Our identification of this branch with putative O-LM cells predicts that expression of these genes should be high in stratum oriens; this prediction was confirmed by examination of in situ hybridization data from the Allen Mouse Brain Atlas (Lein et al., 2007; one example shown in Figure 6a).

**Figure 6.**
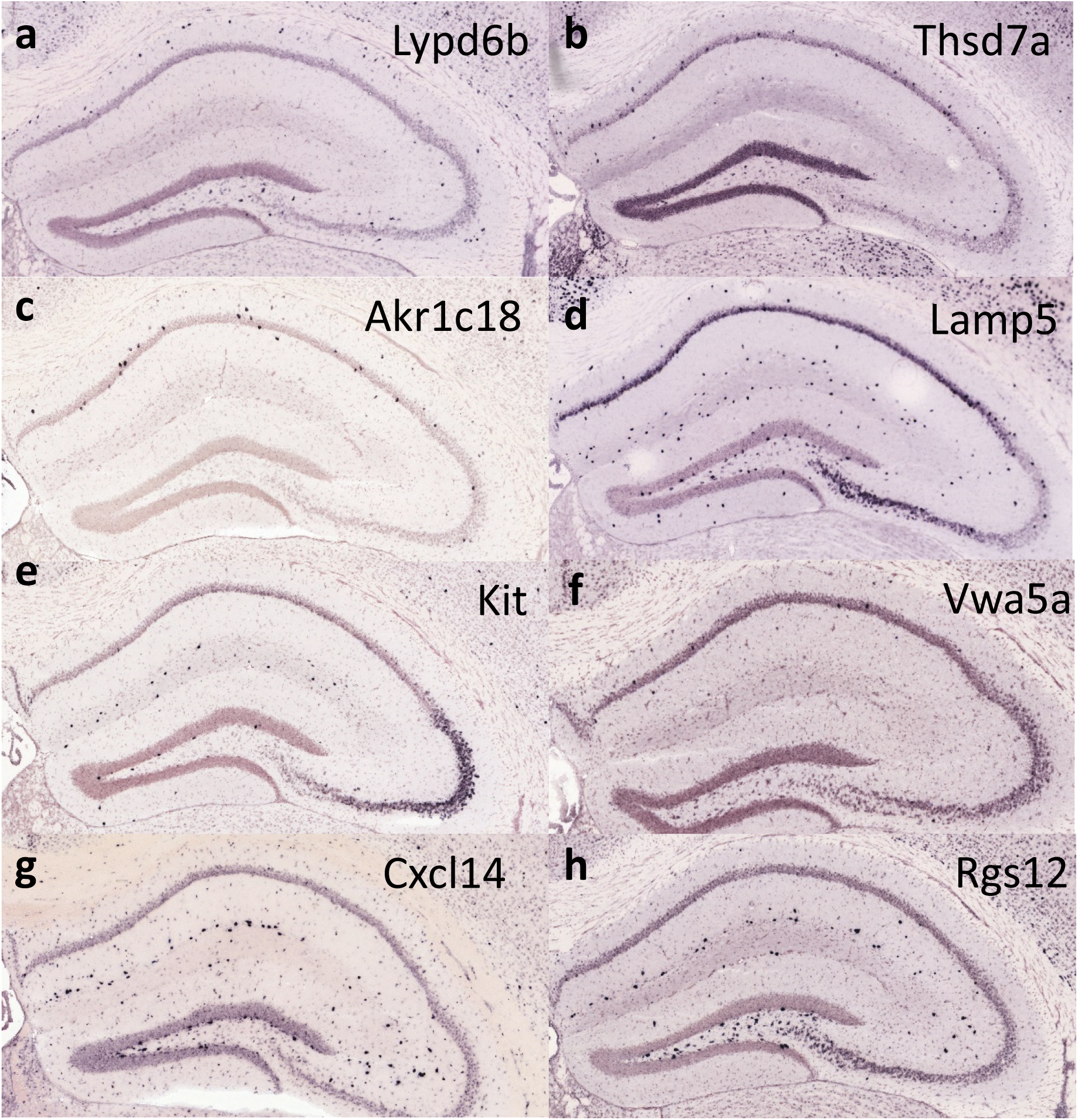
In situ hybridization reveals that novel predicted markers have the expected location within CA1. **(a)**, *Lypd6b*, a predicted marker of O-LM cells shows expression in *stratum oriens*. **(b)**, *Thsd7a*, a predicted marker of basket cells, shows expression in *stratum pyramidale*. **(c)**, *Akr1c18*, a predicted marker of bistratified cells, shows expression in an around *stratum pyramidale*. **(d)**, *Lamp5*, a predicted marker of neurogliaform and ivy cells, shows densest expression in *stratum lacunosum-moleculare* but additional scattered expression around the pyramidal layer. **(e)**, *Kit*, a predicted marker of CGE-derived neurogliaform cells shows expression restricted to *stratum lacunosum-moleculare*. **(f)**, *Vwa5a*, a predicted marker of ivy cells, shows sparse expression around *stratum pyramidale*. **(g,h)**, *Cxcl14* and *Rgs12*, two markers of the novel R2C2 class show expression at the border of *strata radiatum* and *lacunosum-moleculare*. All images from the Allen Mouse Brain Atlas, experiments 73635774, 71924155, 74425542, 70927827, 73520994, 71836830, 74272041, 70613990.

These putative O-LM cells split at the 4^th^ hierarchical level into two further branches (Supplementary Figure 4). This division was less clean than at higher hierarchical levels: there were less genes that distinguished the two branches, and the mutual exclusivity of their expression was less strict. The most notable difference between these branches was their expression of *Htr3a*, a receptor that has been shown to be differentially expressed amongst O-LM cells (Chittajallu et al., 2013). Expression of *Satb1* was high in both the Htr3a-positive and Htr3a-negative branches, consistent with previous reports (Chittajallu et al., 2013); however, we note that expression of *Lhx6* – often considered a marker of MGE-derived interneurons (Cobos et al., 2005, Cobos et al., 2006, Fogarty et al., 2007, Liodis et al., 2007, Du et al., 2008) – was strong in both populations. Thus, while our data are consistent with the existence of subpopulations of O-LM cells that differ in *Htr3a* expression, they do not provide additional evidence in favor of their distinct developmental origins.

Cells in the 3^rd^-level sister branch to the putative O-LM cells expressed at least one of a set of marker genes including *Crh, Pvalb, Kcnip2, Mef2c*, and *Tac1*, and some of them also expressed *Sst, Grm1, Grin3a, Npy*, and *Sstr1*, albeit at lower levels than those of the putative O-LM cells. Further division of this branch revealed a 4^th^ level branch whose expression largely matched that of putative MGE-derived O-LM cells with additional expression of neuropeptides such as *Crh, Tac1*, and *Pnoc*, which we identify as a potentially new class of *Sst* interneuron we term the *Crh/Sst* cell; and a second 4^th^ level branch characterized by strong expression of *Pvalb*.

The *Pvalb*-positive branch contained only 6 cells, which constituted all the strongly *Pvalb* expressing interneurons in the database; this low number likely reflected a low survival rate of *Pvalb* neurons during tissue dissociation. The small number of *Pvalb* cells present meant that further splits of this branch did not reach statistical significance. Nevertheless, the two 5^th^-level sub-branches identified by the GeneTeam algorithm showed gene expression profiles remarkably similar to basket and bistratified cells (Supplementary figure 4). Indeed, while both groups showed strong expression of *Pvalb*, the putative basket cells also showed strong expression of basket-cell associated genes *Gabrd, Mef2c, Chrm2, Kcnc1* (Hajos et al., 1998, Jonas et al., 2004, Ferando and Mody, 2014, Kepecs and Fishell, 2014) but not *Sst*, whereas the putative bistratified cells showed both *Sst* and *Pvalb* expression. Examination of gene expression in these branches suggested *Thsd7a, Pthlh, Nacc2, Plcxd3, Kcnip2* and *Fam19a2* as novel markers for *Pvalb* basket cells, and *Tac1, Akr1c18*, and *Gria3* as potential markers of bistratified cells. Our identification of these branches with putative fast-spiking basket and bistratified cells predicts that expression of these genes should be high in stratum pyramidale; this prediction was again confirmed by *in situ* hybridization data (Figures 6b, c).

## Putative neurogliaform and ivy cells

The other 2^nd^-level subbranch of MGE-like cells was identified as putative neurogliaform (NGF) and ivy cells. Indeed, cells in this branch were positive for *Npy, Nos1, Gabra1* and *Gabrd*, but negative for *Sst, Pvalb, Vip*, and *Cnr1*, as predicted for NGF/ivy cells (Fuentealba et al., 2008a, Olah et al., 2009, Fuentealba et al., 2010, Tricoire et al., 2010). Furthermore, the branch contained cells positive for *Reln*, cells positive for *Kit*, a tyrosine kinase receptor recently identified with NGF cells of isocortex (Miyoshi et al, SFN Abstract 208.06, 2014), and for *Ndnf* which has also been associated with isocortical NGF cells (Tasic et al, SFN Abstract 598.05, 2015). In addition, our database suggested a large number of new markers for the NGF/ivy branch, including *Cacna2d1, Id2, Hapln1, Lamp5*, and *Sema5a*. Consistent with our proposed identification of this branch, these genes were found to be expressed in scattered cell populations both around the pyramidal layer (where ivy cells are found) and in stratum lacunosum-moleculare (where neurogliaform cells are found; Figure 6d).

The putative NGF/ivy branch was split at the 3^rd^ level into branches that differed in their expression of *Htr3a*, as well as several other markers (Supplementary Figure 5). Unlike with putative O-LM cells, the two 3^rd^ level subbranches of putative NGF/ivy cells differed in their expression of *Lhx6*, adding to evidence that they do indeed reflect CGE- and MGE-derived interneurons (Tricoire et al., 2010, Tricoire et al., 2011). The putative CGE-NGF branch was positive for *Nr2f2* as well as *Kit* and *Ndnf*, two markers of CGE-derived isocortical NGF cells, whereas the putative MGE-derived branch did not show expression of these markers. Consistent with previous results from RT-PCR, we detected expression of *Nos1* in the putative MGE-, but not CGE-derived branches (Tricoire et al., 2010). The analysis also suggested *Cplx3* as a potential novel marker of CGE-derived NGF cells that showed little expression in any other class. Consistent with our identification of this branch as neurogliaform (but not ivy) cells, expression of both *Kit* and *Cplx3* was restricted to *stratum lacunosum-moleculare* (Figure 6e); *Ndnf*, which is also expressed in other branches (see below), had a more widespread expression pattern (not shown).

The putative MGE-derived branch split into two 4^th^-level branches (Supplementary figure 5), one of which showed strong expression of *Reln*, while the other showed stronger expression of *Npy*. Based on previous work (Fuentealba et al., 2008a, Fuentealba et al., 2010, Tricoire et al., 2010, Tricoire et al., 2011), we therefore identified these as putative MGE-neurogliaform cells expressing *Reln*, and ivy cells lacking *Reln*. In addition to these genes, we found strong expression of *Vwa5a* in the putative ivy cell branch, suggesting that this gene may be a selective marker for ivy cells. Consistent with our identification of this branch as ivy cells, *Vwa5a* showed expression in scattered cells around the pyramidal layer (Figure 6f).

## Putative *Cck* basket and *Vip* interneurons

Within the CGE-like branch, the algorithm identified two 2^nd^-level branches, which both contained cells expressing *Cck*, but were distinguished by a large number of other markers (Supplementary figure 6). While most of these markers were novel, some were recognizable from previous studies. One 2^nd^-level branch contained cells that expressed *Slc17a8* (a vesicular glutamate transporter also known as *Vglut3* which marks a subset of *Cck* basket cells (Somogyi et al., 2004)), *Vip*, and/or *Tac2*, but did not express *Reln*. We therefore identified this branch as containing putative *Cck*-basket cells, *Vip*-positive interneurons, as well as potentially other cells, and termed it the *Cck/Vip* branch.

The *Vip/Cck* branch split at the 3^rd^ level into two subbranches, one characterized by expression of *Slc17a8*, and the other by *Vip*. As *Slc17a8* has been shown to mark a subset of *Cck* basket cells, we identified this branch with this subset; consistent with this identification, the branch strongly expressed the known *Cck-*basket markers *Cck, Cnr1*, and *Chrm3* (Cea-del Rio et al., 2010); as well as several novel genes including *Krt73, Kctd12, Cadps2, Rgs10*, and *Sema3c*. As expected based on the widespread distribution of *Cck* basket cells, these genes were expressed in scattered populations throughout all layers of CA1 (Allen Mouse Brain Atlas; data not shown). Consistent with immunohistochemical results, only a small fraction of putative *Slc17a8* basket cells showed expression of *Vip* or *Calb1* (Somogyi et al., 2004). This branch appeared to show a further subdivisions into subsets positive for either *Htr3a* or *Tac2*, but these were not analyzed in depth due to the small number of remaining cells. As a whole, the *Slc17a8/Cck* branch strongly expressed mRNA for *Npy*, a molecule that to our knowledge has not yet been histologically examined together with *Cck* or *Slc17a8*.

The other 3^rd^-level branch of *Vip/Cck* cells contained cells strongly expressing *Vip* but lacking *Slc17a8*. This branch split at the 4^th^ level into one branch whose cells also strongly expressed *Cck*, and another which lacked *Cck* with but contained cells expressing *Calb2* or *Penk*. This division closely parallels the previous immunohistochemical division of *Vip* cells into *Cck-*containing basket cells, and interneuron-selective (IN-sel) interneurons, subsets of which have been shown to express *Calb2* and *Penk* (Acsady et al., 1996a, Acsady et al., 1996b, Blasco-Ibanez et al., 1998, Fuentealba et al., 2008b, Tyan et al., 2014). Consistent with this identification, the putative *Vip/Cck* basket cells expressed *Cnr1* as well as several putative novel markers of *Cck* basket cells (Figure 5); the putative IN-sel cells did not strongly express these markers, but a subset of them expressed *Nos1* consistent with previous RT-PCR experiments (Tricoire et al., 2010). This identification allowed us to hypothesize *Hpcal1* and *Qrfpr* as novel markers for IN-sel cells, and *Pcp4* and *Sel1l3* for *Vip/Cck* basket cells. These 4^th^-level subgroups showed evidence for further subdivisions, for example the mutually-exclusive expression of *Calb2* and *Penk* in putative IN-sel cells, but these were not examined in further depth.

## R2C2 cells

The second and final 2^nd^-level branch of the CGE-like interneurons was identified by a number of primarily novel markers. We name this branch the “R2_2” branch, after four of its most characteristic identifying genes: *Rgs12, Reln, Cxcl14*, and *Cpne5*. Examination of laminar expression patterns for the novel markers of this branch suggested that its cells were primarily located at the border of *strata radiatum* and *lacunosum-moleculare* (Figure 6g, h). While this branch contained no single universal and exclusive marker, *Cxcl14* was expressed in all cells of the branch (as well as being expressed at a lower level in *Slc17a8-*positive putative *Cck* basket cells). We were not able to readily identify this branch with cell populations previously described in the literature, and therefore suggest it represents novel molecular classes.

The R2C2 branch split into two 3^rd^-level subbranches. The first of these expressed *Cck* and *Cnr1* at a level exceeded only by the putative *Cck* basket cells, and additionally expressed several other markers associated with them including *Npy, Cadps2*, *Car2, Chrm3, Kctd12, Krt73, Rgs10, Sema3c*, and *Snca*, but did not express *Slc17a8*, and only rarely *Vip* (Figure 5; Supplementary Figure 6). Based on their inferred location at the *strata radiatum/lacunosum-moleculare* border, we hypothesize that this branch might correspond to a population of *Cck*-positive non-basket cells, such as perforant-path associated cells (Klausberger et al., 2005). Consistent with this identification, we note that many cells in the branch expressed *Calb1*, a molecule that has been detected in a subset of those *Cck-*positive neurons expressing neither *Slc17a8* nor *Vip*, and likely correspond to interneurons targeting pyramidal cell dendrites (Cope et al., 2002, Somogyi et al., 2004, Klausberger et al., 2005). As with putative O-LM and NGF cells, this branch showed an additional 4^th^-level division into *Htr3a*-positive and *Htr3a*-negative subbranches, which showed few other major differences in expression profile (not shown).

The other 3^rd^-level branch of *Cxcl14* cells was negative for *Npy* and most other novel markers associated with putative *Cck* basket cells, but contained cells expressing new markers including *Igfbp5* and *Ndnf*, as well as the strongest expression of *Calb1* of all branches. These cells expressed *Nr2f2*, and we hypothesize that they might correspond to a population of *Nr2f2*-positive cells in *str. radiatum* and *lacunosum-moleculare* that were not marked by other known molecules (Fuentealba et al., 2010). This branch split into two 4^th^ level subbranches (Figure 4; Supplementary figure 6). The first of these was characterized by expression of *Tnfaip8l3* as well as additional markers including *Slc7a14, Stxbp6* and *Ndnf* (note that while *Ndnf* is also found in *Kit*-positive putative CGE-neurogliaform cells, we do not hypothesize the present branch correspond to neurogliaform cells due their weak expression of *Gabrd* and general molecular dissimilarity to the *Kit-*expressing population). This branch expressed *Cck* mRNA at low levels; however, we note that this does not necessarily imply expression of protein, as *Cck* expression in these cells was no higher than in putative *Pvalb* basket cells (c.f. Tricoire et al., 2011). This branch also showed the strongest expression of *Npy2r* we observed in the database; consistent with previous immunohistochemical analysis (Stanic et al., 2011), the cells expressing this receptor showed no expression of *Npy* itself.

The final 4^th^-level subbranch of *Igfbp5* cells was identified by strong expression of *Ntng1* but weak expression of *Cck*. This class contained some cells positive for *Vip* and *Penk*, and we therefore hypothesize it represents a second set of IN-sel interneurons located at the R-LM border.

## Testing predictions of the classification with immunohistochemistry and *in situ* hybridization

The classification scheme we derived makes a large number of predictions for the combinatorial expression patterns of familiar and novel molecular markers in distinct CA1 interneuron types. We next set out to test some of these predictions using traditional methods of molecular histology.

A strong prediction of our classification was the expression of *Npy* in two branches that also expressed *Cck* (the *Slc17a8/Cck* branch of *Calb1-*negative putative basket cells; and the *Calb1*-positive, *Slc17a8-*negative R2C2/*Cck* branch). This was unexpected, as NPY (at least at the protein level) has instead been traditionally associated with SST-expressing neurons and ivy/neurogliaform cells (Fuentealba et al., 2008a, Katona et al., 2014). Nevertheless, no studies to our knowledge have yet examined immunohistochemically whether the neuropeptides NPY and CCK can be colocalized in the same interneurons. We therefore tested this by double immunohistochemistry in *strata radiatum* and *lacunosum-moleculare* (Fig. 7a-b, n=3 mice). Consistent with the model’s prediction, 119 out of 162 (74±6%) of the cells immunopositive for pro-CCK were also positive for NPY (an additional 73 cells were positive for NPY only, which according to our classification scheme should represent primarily neurogliaform cells). A subset (n=176) of NPY and/or pro-CCK immunopositive neurons were further tested for CALB1 (calbindin) in triple immunoreactions. As expected, nearly all CALB1-positive neurons were pro-CCK-positive (89±2%), and CALB1 immunoreactivity was seen in a subset of the cells containing both pro-CCK and NPY (27±3%). Additional triple immunohistochemistry for NPY, pro-CCK and SLC17A8 (VGLUT3) revealed triple positive cells in *stratum radiatum* and particularly at the border with *lacunosum moleculare* (Figure 7b). Due to the low level of somatic immunoreactivity of SLC17A8 (which as a vesicular transporter is primarily trafficked to axon terminals), we could not count these cells reliably; however of the cells that were unambiguously immunopositive for SLC17A8, in a majority we detected NPY, consistent with the model predictions. Additional analysis combining double *in situ* hybridization for *Slc17a8* and *Npy* with immunohistochemistry for pro-CCK (Figure 7c, n=3 mice) confirmed that the great majority of *Slc17a8*-expressing cells were also positive for *Npy* and pro-CCK (84±3%). As predicted by our classification scheme, the converse was not true: a substantial population of *Npy*/pro-CCK double-positive cells (57±7% of the total) did not show detectable *Slc17a8*, which according to our model should primarily consist of R2C2/*Cck* cells.

**Fig 7.**
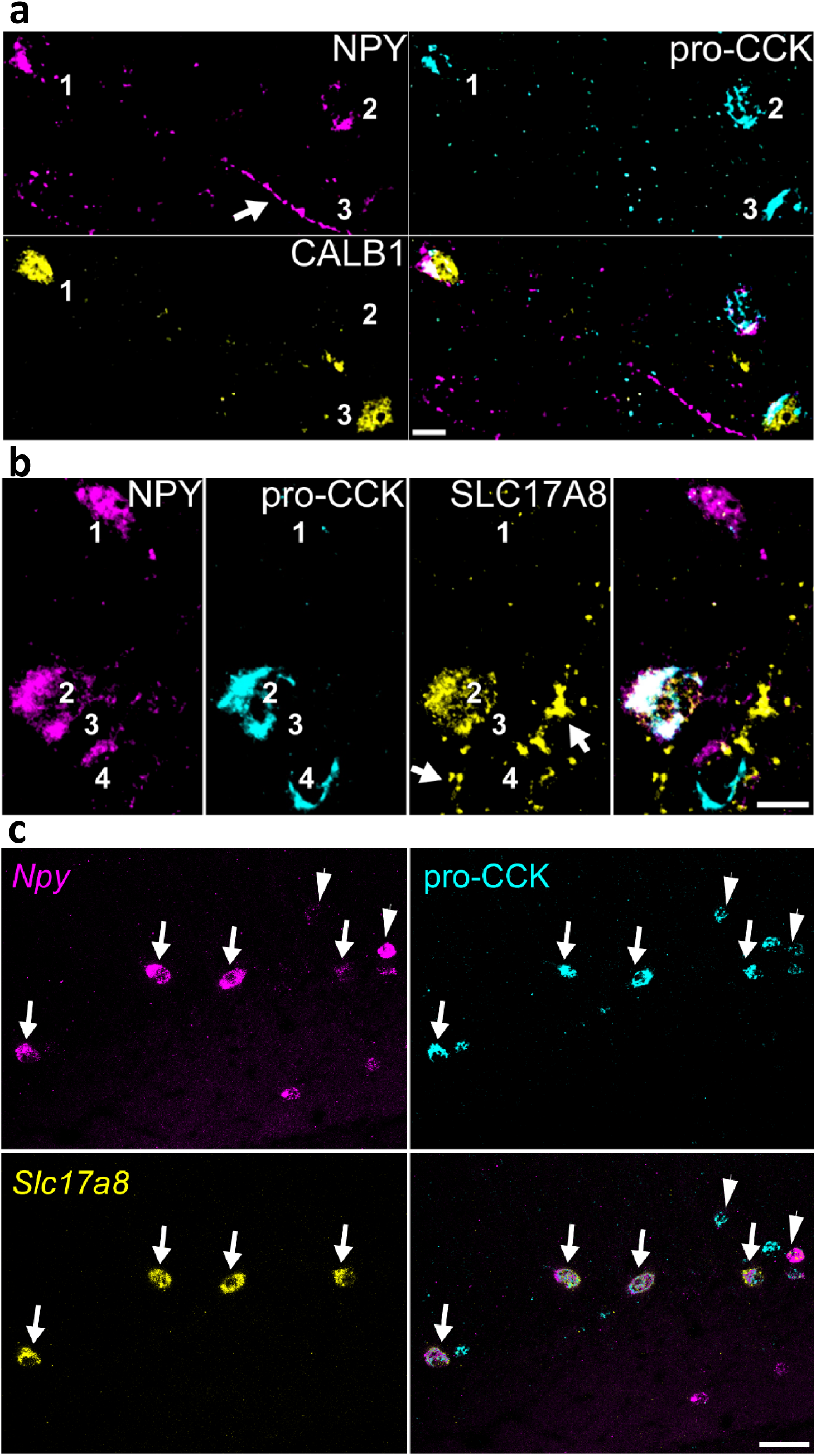
Confirmation of predicted co-localisation of NPY and pro-CCK. **(a)**, Interneurons at the border between strata radiatum and lacunosum moleculare are immunopositive for both NPY and pro-CCK (cells 1 and 2), one of which (cell 1) is also immunopositive for CALB1. A third neuron is positive only for pro-CCK and CALB1 (cell 3). **(b)**, Interneurons at the border of strata radiatum and lacunosum moleculare are immunopositive for NPY (cells 1–3), pro-CCK (cells 2 and 4) and SLC17A8 (VGLUT3, cell 2). Note SLC17A8-positive terminals targeting unlabelled cells (arrows). (a,b), Both NPY and pro-CCK are detected in the Golgi apparatus and endoplasmic reticulum surrounding cell nuclei, in addition, some axons are also immunopositive for NPY (see arrow in (a); average intensity projections, z stacks, height 6.3 µm and 10.4 µm, respectively). **(c)**, Combined double *in situ* hybridization and immunohistochemistry shows that nearly all *Slc17a8*-expressing cells also express *Npy* and are immunopositive for pro-CCK (arrows), but some *Npy/*pro-CCK cells do not express *Slc17a8* (arrowheads). Scale bars: 10 µm (a,b), 50 µm (c).

Our scRNA-seq data suggested the existence of a new class of hippocampal interneurons (the R2C2 cells) located at the *radiatum*/*lacunosum* border (R-LM), and characterized by expression of *Cxcl14* as well as other novel markers. According to the classification scheme we derived, these cells should be mostly positive for *Reln;* negative for *Pvalb, Sst* and other markers of MGE-derived populations; should contain subpopulations positive for *Cck, Calb1* and *Npy;* and should be distinct from neurogliaform cells. To test these predictions, we performed *in situ* hybridization for *Cxcl14* simultaneously with in situ hybridization or immunohistochemistry to detect *Reln, Npy*, CALB1, CCK, PVALB, *Sst* as well as *Nos1* and *Kit* as markers of neurogliaform interneurons (n=3 mice; italics denote molecules detected by *in situ* hybridization, capitals by immunohistochemistry). In addition, we combined fluorescent *in situ* hybridization for *Cxcl14* with immunohistochemistry for YFP in *Lhx6-Cre/R26R-YFP* mice, which allows identification of developmental origin by marking MGE-derived interneurons (Fogarty et al., 2007).

The results of these experiments were consistent with our hypotheses. We found that within CA1, *Cxcl14-*expressing cells were primarily located at the R-LM border (71±3%), although a subpopulation of cells were also found other layers (possibly corresponding to *Slc17a8*-positive basket cells, which also express *Cxcl14*). We found no overlap of *Cxcl14* with YFP in the *Lhx6-Cre/R26R-YFP* mouse, confirming the CGE origin of *Cxcl14* expressing neurons (Figure 8a); consistent with this finding, no overlap was seen with *Sst* or *Pvalb* (data not shown). The majority of *Cxcl14*-positive cells expressed *Reln* (72±4%), although a smaller fraction of Reln-expressing neurons were *Cxcl14* positive (42±9%), with substantial populations of *Reln*+/*Cxcl14-*cells in *strata oriens* and *lacunosum-moleculare* likely representing O-LM and neurogliaform cells, respectively (Fig 8b). Indeed, although less than half of *Reln* cells were located at the R-LM border (44±1%), the great majority of *Reln+/Cxcl14+* cells were found there (88±6%), consistent with the expected location of R2C2 neurons. Also consistent with the model, a large fraction of the *Cxcl14* population were immunopositive for pro-CCK (62±6%; Fig. 8c), while substantial minorities were positive for CALB1 (29±2%; Fig 8d) or *Npy* (25±5%; Fig 8e). However, we observed essentially no overlap of *Cxcl14* with *Nos1* or *Kit* (0 of 209 and 1 of 264 cells respectively, from all mice), suggesting that these neurons are distinct from the neurogliaform population. The results of this double labeling analysis are therefore consistent with the predictions of our classification model.

**Fig 8.**
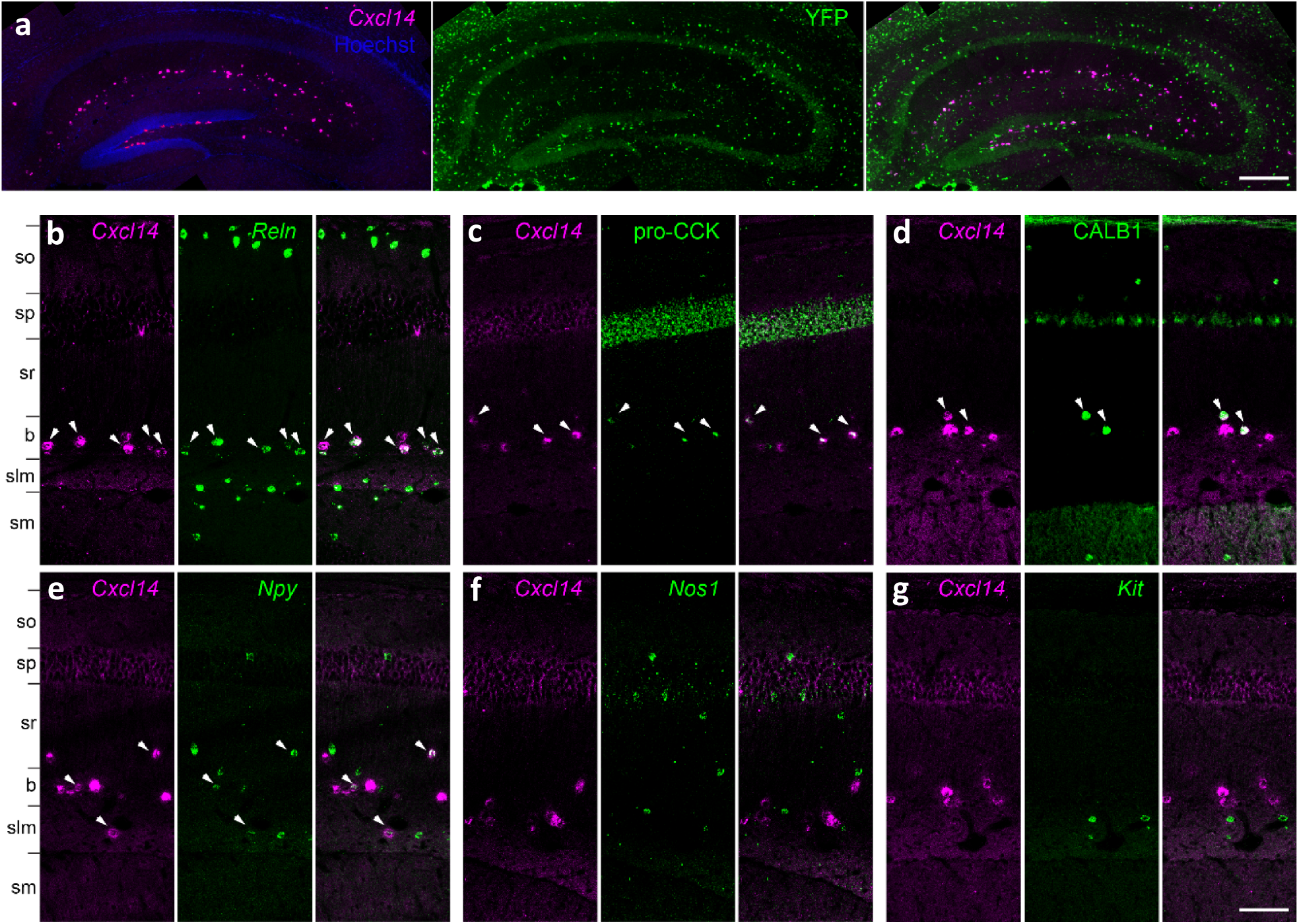
Analysis of *Cxcl14* co-expression patterns confirms predicted properties of R2C2 cells. **(a)**, *Cxcl14-*expressing cells are CGE-derived: in situ hybridization for *Cxcl14* combined with immunohistochemistry for YFP in the *Lhx6-Cre/R26R-YFP* mouse yields no double labelling. **(b)**, Double *in situ* hybridization for *Cxcl14* and *Reln* marks a population of neurons located primarily at the border of *strata radiatum* and *lacunosum-moleculare*, as predicted for R2C2 cells. Note *Reln* expression without *Cxcl14* in *strata oriens* and *lacunosum-moleculare*, likely reflecting O-LM and neurogliaform cells. **(c-e)**, Subsets of the *Cxcl14-*positive neurons are positive for pro-CCK or CALB1 (*in situ* hybridization plus immunohistochemistry), or *Npy* (double *in situ* hybridization). **(f, g)** No overlap was seen of *Cxcl14* with *Nos1* or *Kit*. In all panels, arrowheads indicate double-expressing neurons. Layer abbreviations: so, *stratum oriens;* sp, *stratum pyramidale;* sr, *stratum radiatum;* b, R-LM border region; slm*, stratum lacunosum-moleculare;* sm, *stratum moleculare* of the dentate gyrus. Scale bars: 200 µm (a), 100 µm (b-g).

## Discussion

We have applied a novel analysis algorithm to scRNA-seq data, to derive a classification of CA1 interneurons into a 5-level hierarchy with 14 identified classes. The expression patterns of most of these classes bear a striking resemblance to classes defined by previous immunohistochemical work, allowing us to putatively identify the cell types represented by most branches, down to fine levels such as ivy cells, and *Cck* basket cells positive for *Slc17a8* or *Vip*. The fact that our methods accurately predicted known interneuron classes provides confidence in the novel classes produced by our classification scheme; in addition, we directly confirmed several new predictions of the scheme using *in situ* hybridization and immunohistochemistry.

The largest putative novel interneuron class we identified (the R2C2 branch) was characterized by combinatorial expression of novel markers including *Rgs12, Reln, Cxcl14*, and *Cpne5*, with some cells within this class also expressing previously studied molecules such as *Calb1, Npy, Cck, Tnfaip8l3*, and *Nr2f2*. The laminar expression profile of markers for this class suggests a concentration around the border of *strata radiatum* and *lacunosum-moleculare*, a result we confirmed using *in situ* hybridization, as well as by confirming the predicted patterns of overlap for additional molecular markers. Cells in this spatial location have been relatively less well studied in the literature, but contain several populations including basket cells innervating the somata of CA1 pyramidal neurons; Schaffer collateral-associated, apical-dendrite-targeting, and perforant-path associated neurons innervating pyramidal cell dendrites; neurogliaform interneurons; interneuron-selective interneurons; as well as long-range GABAergic projection cells with targets in retrosplenial cortex (Vida et al., 1998, Klausberger et al., 2005, Jinno et al., 2007, Miyashita and Rockland, 2007, Fuentealba et al., 2008b, Fuentealba et al., 2010, Melzer et al., 2012, Kitamura et al., 2014). It is likely that the R2C2 cells comprise only a subset of these neurons, rather than the entire population. We were able to divide the R2C2 cells into three subgroups, which we propose correspond to a set of dendrite-targeting *Cck* interneurons; a set of interneuron-selective neurons; and a set of *Tnfaip8l3*-positive neurons which express few other known markers, and whose connectional relationships remain to be identified.

The classification we have derived here is likely to underestimate the true complexity of CA1 interneuron types. Indeed, while the major molecular classes described in the literature have been identified in the current classification scheme, there exist additional specific classes which have not been found. For example, interneurons projecting from CA1 to distal targets such as the medial septum or subiculum express *Sst* in combination with other markers such as *Calb1, Calb2*, and *Chrm2* (Jinno, 2009); while we found individual cells in the database matching these molecular profiles, their small numbers resulted in grouping together with putative O-LM cells. Subdivisions of basket, bistratified, and axo-axonic neurons have been reported that differ in their somadendritic laminar organization and spike timing relative to LFP oscillations (Varga et al., 2014); while we observed heterogeneity in the *Pvalb*-positive population, the present sample size was too small to reliably identify further subgroups. Interestingly, a class of hippocampal interneuron that projects to the retrosplenial cortex, but is negative for most classical molecular markers, is located at the border of *strata radiatum* and *lacunosum-moleculare* (Miyashita and Rockland, 2007); we speculate that this class might correspond to one of the novel R2C2 classes we have identified.

Immunohistochemical analysis has suggested that CA1 interneuron types are identified not by single marker genes, but by combinatorial expression patterns (Somogyi, 2010). However such studies cannot exclude the possibility that unique identifiers exist, but have not yet been tested. Because scRNA-seq scans the entire genome it offers a greater opportunity to find genes uniquely identifying cell classes. We found such genes only rarely, for example *Vwa5a* as a putative unique identifier of ivy cells, and *Cplx3* and *Kit* as putative unique identifiers of CGE-derived neurogliaform cells. Despite this lack of unique identifiers, the ability of scRNA-seq to characterize multiple genes simultaneously allowed us to predict novel combinatorial relationships between classical marker genes (such as the expression of *Npy* in *Cck* interneurons), and to identify a large number of candidate marker genes for familiar and novel classes, such as *Cxcl14, Lamp5, Rgs12, Igfbp5, Crhbp, Ntng1* and *Ndnf*. It will be possible to test more of these markers in the predicted combinatorial expression patterns in future systematic studies.

The novel molecular markers we identified have been associated with highly diverse biological functions including synaptic or vesicular function (e.g. *Cacna2d1*, *Rab3b, Cpne5, Cplx3, Nxph1*); intercellular adhesion and recognition (e.g. *Cdh13, Sema3c, Ndnf, Wnt5a, Ntng1, Cxcl14*); peptides, receptors and signaling (*Crh, Crhbp, Igf1, Igfbp5, Rgs12, Rgs17, Gucy1a3)*; extracellular matrix proteins (*Hapln1, Col19a1, Col25a1*); and even a molecule best known as a constituent of hair (*Krt73*). One surprise was the relatively small number of transcription factors identified; it may be that the relatively low expression levels of these molecules precluded their detection by our methods.

Do interneurons truly divide into discrete classes, or are they points along a continuum (Parra et al., 1998, Markram et al., 2004)? Our data suggest the existence of discrete classes divided by expression of large mutually-exclusive gene sets, but also support the existence of a continuum of expression levels for some molecules within each of these classes (Supplementary Figure 7). RNA transcription levels are dynamically modulated (Kaern et al., 2005), and this modulation can correlate with behavior. The expression level of *Pvalb* in CA3 basket cells, for example, is modulated by neuronal activity and learning (Donato et al., 2013). Nevertheless, this modulation does not cause *Pvalb*-negative neurons to become *Pvalb*-positive, nor does it drive *Pvalb* expression in basket cells all the way to zero. Similarly, the expression of the transcription factor *Er81* and potassium channel *Kcna1* in isocortical basket cells is modulated by activity (Dehorter et al., 2015), but these modulations in activity do not drive *Er81* expression fully to zero. We therefore hypothesize that dynamic modulation of expression can change a cell’s expression levels within the continuum defined by a single cell class, but that cells will rarely if ever “jump” from one class to another (Supplementary Figure 7). Indeed, the long-term stability of interneuron classes is supported by the non-overlapping and mutually exclusive placement of synapses from individual interneurons on distinct subcellular domains of pyramidal cells, together with their exquisitely cell-type dependent temporal discharge patterns (Katona et al., 2014, Somogyi et al., 2014, Varga et al., 2014).

In summary, we have used single-cell sequencing data to derive a hierarchical classification of CA1 interneurons. This classification confirmed many cell categories previously derived by immunohistochemistry and connectivity, including at deep levels of the classification tree, and suggested several new cell classes together with predicted expression patterns for a large number of new and familiar molecular markers in each class. This analysis confirms the stunning diversity of CA1 interneuron types, and raises the possibility that a similar level of complexity in GABAergic neuronal populations occurs throughout cortex, and indeed throughout the brain. This fine diversity of interneuron classes likely underpins an exquisite regulation of hippocampal information processing and plasticity.

## Experimental Procedures

### Gene team algorithm

The gene expression data takes the form of a matrix *x_cg_* of non-negative integers, each entry of which represents the number of RNA molecules of gene *g* detected in cell *c*. We write **x***_c_* for the *N_genes_* -dimensional expression vector corresponding to cell *c*. Methods by which the data were collected are described in (Zeisel et al., 2015), and the data are available online at http://linnarssonlab.org/cortex/. (Note that while the analysis of Zeisel et al started from a selected subset of 5000 genes, all 19,972 were used here.)

The Gene Team algorithm is a divisive hierarchical clustering method. The algorithm operates recursively: it begins by dividing the dividing the full set of cells into two branches; then the algorithm is re-run on each of these branches; and so on, providing a multi-level hierarchical classification.

The algorithm splits a set of cells into two branches by finding two “teams” of genes, such that each cell strongly expresses at least one gene from one team, but expresses none of the genes from the other team. More specifically, the two teams are represented by *N_genes_*-dimensional weight vectors **w_1_** and **w_2_**. We define the “team scores” for cell *c* as *X_c_* = **x_c_ · w_1_** and *Y_c_* = **x_c_ · w_2_**. The weight vectors **w_1_** and **w_2_** are constrained to be non-negative, and a penalty function (described below) ensures that most entries will be zero; thus these vectors define two discrete teams (i.e. subsets) of genes.

The extent to which expression of the two gene teams is mutually-exclusive is captured by an objective function, illustrated in figure 2a, and defined by the formula

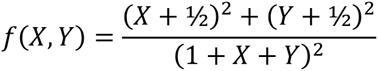

A straightforward calculation shows that ½ ≤ *f*(*X*, *Y*) < 1, and that the constant contours of *f*(*X*, *Y*) = *F* are given by:

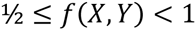

Thus, fixed values of *F* correspond to straight lines through the point (-½, -½), with a slope that gets further above or below 45° as the value of *F* increases. Because *X_c_* and *Y_c_* are both constrained to be non-negative, obtaining a large value of F therefore requires that one of *X_c_* or *Y_c_* be large, and the other close to zero. When *X_c_* and *Y_c_* are equal (including when they are both equal to zero), score takes its minimal value. Thus, a high score will be obtained if a cell strongly expresses at least one gene from one team, but no genes from the other team.

By maximizing the sum of *f*(*X_c_*, *Y_c_*) over all cells, the algorithm therefore ensures cells are divided into two groups, of which one lies close to the x-axis, and the other lies close to the y-axis. This gives rise to the characteristic “L-shaped” plots, while also ensuring as few points as possible are near the origin, enabling accurate classification of all cells.

Optimization of *f*(*X*, *Y*) alone is insufficient to classify cells, for two reasons. First, a higher score could always be obtained by making the weights **w**_1_ and **w**_2_ larger, regardless of separation quality. To solve this, we add a penalty term equal to 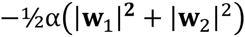, where *α* is a parameter equal to 0.05 in the current study. Second, a large value of ∑*_c_ f*(*X_c_*, *Y_c_*), could be obtained even if all cells were assigned to a single class. To avoid this possibility, we added a second penalty term. Defining the function *g*(*X*, *Y*) as

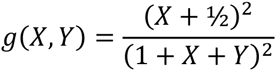

we note that *f*(*X_c_*, *Y_c_*) = *g*(*X_c_*, *Y_c_*)+*g*(*Y_c_*, *Xc*), with the two terms representing the contribution of points near the *x* and *y* axes, respectively. We define a second penalty term 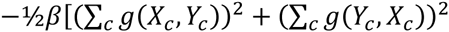, where 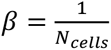. This term may be understood as an *L*_2_ penalty on a 2-dimensional vector approximately equal to the number of cells in each class, and will therefore favor division of the cells into close-to-equal size classes.

The full objective function is thus given by:

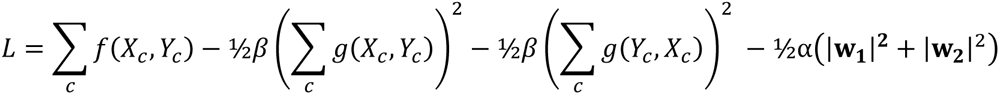

The objective function *L* is optimized over the weight vectors **w_1_** and **w_2_** numerically, subject to the constraints that all elements are non-negative. Because optimization in the full *N_genes_*-dimensional space would be impractical, the search is performed only on the 100 genes for which the cells being classified show most bimodal expression, i.e. genes whose expression histogram has a large peak at zero as well as another peak at positive mean [see e.g. (Shalek et al., 2014)]. As the expression levels are integers, we assess the bimodality of each gene’s distribution as the excess fraction of cells expressing zero copies of that gene, compared to what would be expected under a negative binomial distribution of reasonable skewness. Specifically, a negative binomial distribution is fitted to each gene’s expression histogram by maximum likelihood subject to the constraint that .2 < *p* < .99, and that gene’s bimodality index is computed as the probability of observing zero in the data minus the prediction of the fitted negative binomial distribution. Genes with consistently weak expression are excluded from the search (specifically, genes for which less than 5 cells expressed more than 4 copies of the gene).

To perform the constrained optimization, we used sequential quadratic programming (implemented in MITLAB’s optimization toolbox), using an analytically-computed derivative function to speed up search time. Because the objective function is non-convex, searches were initiated from 100 start points, corresponding to **w_1_** being each unit vector in the 100-dimensional search space, and **w_2_** = 0. Typically, this led to a small number of local optima being found repeatedly, and the local optimum with highest objective function was kept.

To assess the statistical significance of each branch division, we created a null distribution by optimizing the same objective function on an ensemble of random gene expression matrices generated for the cells in the parent branch. Because mean expression levels differ widely across genes, and across cells, we used Patefield’s algorithm (Patefield, 1981; MATLAB implementation by John Burkardt, http://people.sc.fsu.edu/~jburkardt/m_src/asa159/asa159.html) to generate 50 random integer matrices whose row and column marginals match those of the original expression data. The optimal scores for these 50 random matrices were fit by an extreme value (Gumbel) distribution, and statistical significance was computed from the percentile value of the original score within this distribution.

Recursive subdivision of branches continued until splits were no longer statistically significant (p<10^−4^). Importantly, however, the fact that the algorithm could not find a statistically significant split of a branch does not indicate that this branch is a homogeneous cell type; this could occur simply because this branch contained too few cells. Indeed, the algorithm suggested a 5^th^ level division that matched markers expected from previous molecular analyses; this is indicated by dashed lines in Figure 4. In a small number of cases, splits were found corresponding to sex-specific genes (e.g. *Xist, Tsix, Ddx3y, Eif2s3y*), indicating that the division found by the algorithm reflected the gender of the host animal; in these cases, this local optimum was skipped and the second-best fit used. In one case (the division of the *Vip* branch into *Vip/Cck* and IN-sel branches), an additional local optimum was skipped as it reflected a poor separation. For the classification described in the current manuscript, the algorithm was run on a dataset in which a small set of genes likely to represent contamination by glia or pyramidal cells was removed (*Plp1, Crym, Epha4, Sv2b, Neurod6, Prkcb*)

## Relative expression index

To enable rapid identification of genes differentially expressed between two branches, we developed a visualization method where gene names were colored according to their expression levels compared to the entire population (Supplementary Figures 3–6). Specifically, gene names were colored according to an expression index

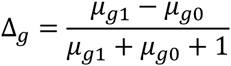

Here *μ_g_*_1_ represents the average expression of gene *g* over all cells in the branch of interest, and *μg*_0_ represents its average expression in all other cells of the database. (To avoid overweighting of outliers, averages were estimated as 1/3 trimmed means, i.e. the mean of the central third of the distribution). The addition of 1 to the denominator served to regularize the index, reducing its value for genes of overall weak expression. The index approaches the value +1 for genes expressed strongly and exclusively in the branch of interest, and -1 for genes that are not expressed in this branch, but are expressed strongly in at least a subset cells outside the branch. Genes whose expression is on average equal inside and outside the branch while have an expression index of 0, and be represented by white colors

## Immunohistochemical analysis and cell counting

Three adult (20 weeks old) male C57BL/6J mice (Charles River, Oxford, UK) were perfusion fixed following anesthesia as described (Unal et al., 2015). Tissue preparation for immunofluorescence (Katona et al., 2014) and analysis using wide-field epifluorescence microscopy (Somogyi et al., 2004) were performed as described previously. The following primary antibodies were used: anti-calbindin (goat, Fronteir Inst, Af104)); anti-pro-CCK (rabbit, 1:2000, Somogyi et al., 2004); anti-NPY (mouse, 1:5000, Abcam, #ab112473); anti-VGLUT3 (guinea pig, Somogyi et al 2004). For cell counting, image stacks (212 x 212 µm area; 512 x 512 pixels; z stack height on average 12 µm) were acquired using LSM 710/AxioImager.Z1 (Carl Zeiss) laser scanning confocal microscope equipped with Plan-Apochromat 40x/1.3 Oil DIC M27 objective and controlled using ZEN (2008 v5.0 Black, Carl Zeiss). In a second set of sections, images were taken using Leitz DM RB (Leica) epifluorescence microscope equipped with PL Fluotar 40x/0.5 objective. Counting was performed either using ImageJ (v1.50b, Cell Counter plugin) on the confocal image stacks or OPENLAB software for the epifluorescence documentation. Numbers were pooled from two separate reactions testing for a given combination of primary antibodies (n=3 mice each reaction, 2–3 sections each mouse). Image processing was performed using ZEN (2012 Blue, Carl Zeiss) and Photoshop (CS5, Adobe). Percentages of double-labelled cells are given as the mean over all mice, ± standard error.

## *In situ* hybridization and cell counting

Wild type (C57BL/6/CBA) male adult (P30) mice and *Lhx6-Cre^Tg^* transgenic mice were perfusion-fixed as previously described (Rubin et al., 2010), followed by immersion fixation overnight in 4% paraformaldehyde. Fixed samples were cryoprotected by overnight immersion in 20% sucrose, embedded in optimal cutting temperature (OCT) compound (Tissue Tek, Raymond Lamb Ltd Medical Supplies, Eastbourne, UK) and frozen on dry ice. 30 µm cryosections were collected in DEPC-treated PBS and double *in situ* hybridization was carried out as described (Rubin et al., 2010). RNA mouse probes were transcribed from the following I.M.A.G.E. (Integrated Molecular Analysis of Genomes and their Expression) clones: *Cxcl14* (ID 30358501), *Sst* (ID 4218815), *Scl17a8* (ID 5367275), or as described elsewhere (Magno et al., 2012). Probes used included either a *Cxcl14*-(digoxgenin)DIG RNA probe in combination with *Reln*-(fluorescein)FITC, *Npy*-FITC or *Sst*-FITC probes, or a *Cxcl14*-FITC probe with *Nos1*-DIG, *Kit*-DIG, *Scl17a8*-DIG, or *Pvalb*-DIG probes. DIG-labelled probes were detected with an anti-DIG-alkaline phosphatase (AP)-conjugated antibody (1:1000, Sigma-Aldrich, Dorset, UK) followed by application of a Fast Red (Sigma-Aldrich, Dorset, UK) substrate. The first reaction was stopped by washing 3 × 10 min in PBS, and the sections were incubated with an anti-FITC-Peroxidase (POD)-conjugated antibody (1:500, Sigma-Aldrich,Dorset, UK) overnight. The POD signal was developed by incubating the sections with Tyramide-FITC:amplification buffer (1:100, TSA™-Plus, Perkin Elmer, Seer Green, UK) for 10 minutes, at room temperature. For immunohistochemistry after *in situ* hybridization the following antibodies were used: anti-Calbindin (rabbit, 1:1000, Swant, Bellinzona, Switzerland); anti-pro-CCK (rabbit, 1:2000, Somogyi et al., 2004); anti-GFP (chicken, 1:500, Aves Labs Inc., Tigard, OR, US). All sections were counterstained with Hoechst 33258 dye (Sigma-Aldrich, Dorset, UK, 1000-fold dilution) and mounted with Dako Fluorescence Mounting Medium (DAKO UK Ltd, Cambridgeshire, UK).

For cell counts, images (at least two sections per mouse) were acquired on an epifluorescence miscroscope (Zeiss) with a 10x objective. Several images spanning the entire hippocampal CA1 were stitched using Microsoft Image Composite Editor. Cells were counted manually in the CA1 area including *strata radiatum* and *lacunosum-moleculare*, and in a subregion spanning 100 µm across the border between *strata radiatum* and *lacunosum moleculare*, where most *Cxcl14*-positive cells are located. Confocal images (z stack height on average 25 µm, 2 *µ*m spacing) were taken on a Leica confocal microscope under a 10x objective and processed for contrast and brightness enhancement with Photoshop (CS5, Adobe). A final composite was generated in Adobe Illustrator (CS5, Adobe). Percentages of double-labelled cells are given as the mean over all mice, ± standard error.

## Acknowledgements

We thank M. Carandini, A. Joshi, G. Unal, T. Viney, and B. Bekkouche for discussions and comments on the manuscript. This work was supported by the Wellcome Trust (108726 to K.D.H., N.K., P.S., S.L., and J.H-L), Medical Research Council (P.S.) European Research Council (261063 to S.L.), Swedish Research Council (STARGET to S.L., and 2014-3863 to J. H-L), Karolinska Institutet (BRECT to A.M-M), StratNeuro (J. H-L), and European Union FP7/Marie Curie Actions (322304 to J.H-L).

---

**Supplementary Figure 1, associated with Figure 3.**
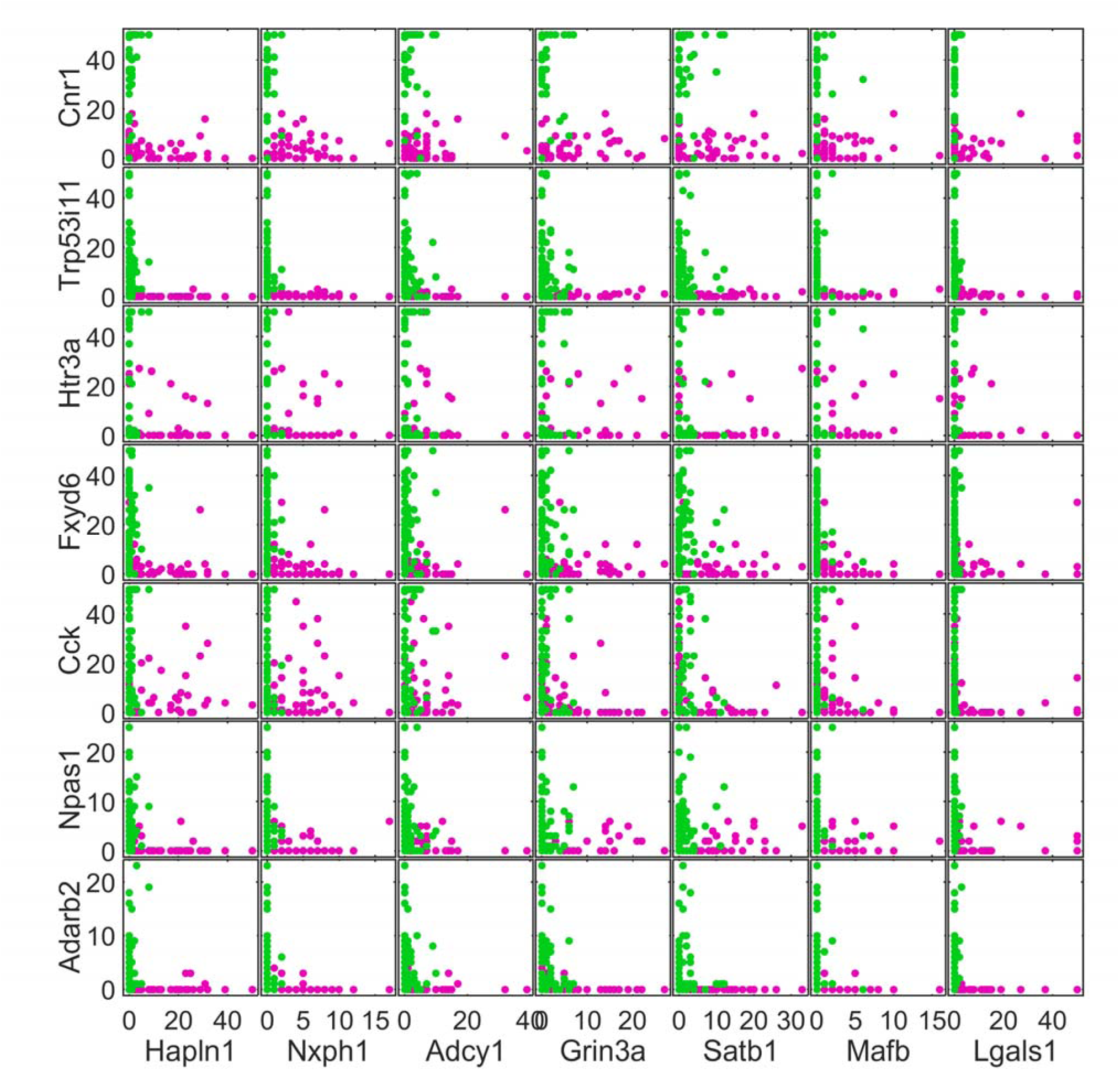
Scatterplot matrix showing expression levels of a subset of additional genes whose expression differs between the MGE‐like and CGE‐like branches. Expression levels for each gene have been clipped to a maximum of 50 molecules/cell to aid visibility. Magenta and green dots represent cells belonging to the MGE‐like and CGE‐like branches, respectively.

**Supplementary Figure 2, associated with Figure 4.**
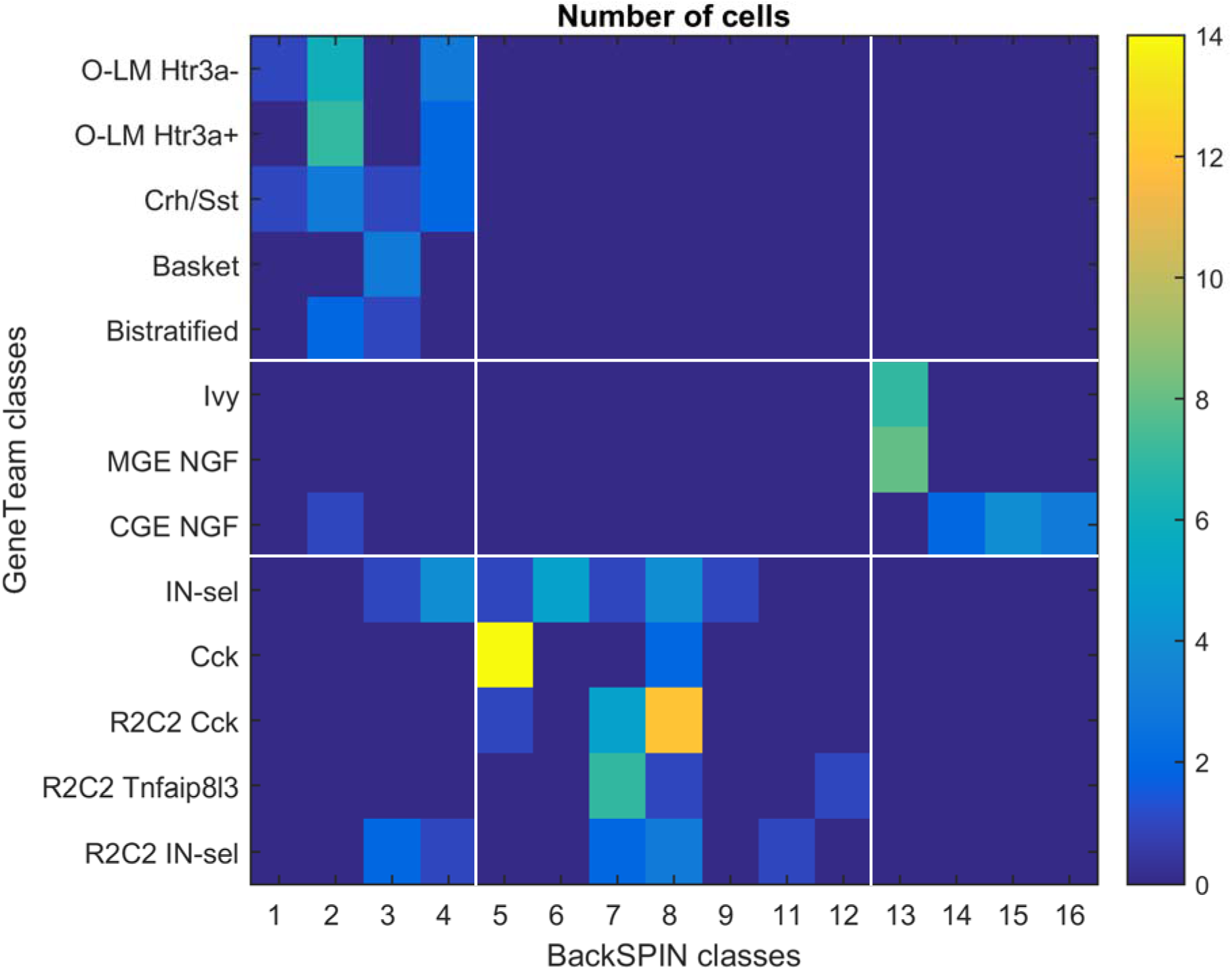
Confusion matrix comparing the classifications of the GeneTeam and BackSPIN algorithms. The classifications agree at a top hierarchical level, with GeneTeam’s *Sst/Pvalb* classes matching BackSPIN’s classes 1‐4; GeneTeam’s *Cck/Vip* and *Rgs12* classes matching BackSPIN’s classes 5‐12; and GeneTeam’s NGF and Ivy cells matching BackSPIN’s classes 13‐16; these top‐level correspondences are marked by white lines. Nevertheless, the results of the two algorithms were not identical at deeper classification levels (note that in addition to the different algorithm, BackSPIN classes were derived from a data set containing both hippocampal and isocortical interneurons).

**Supplementary Figure 3, associated with Figure 4.**
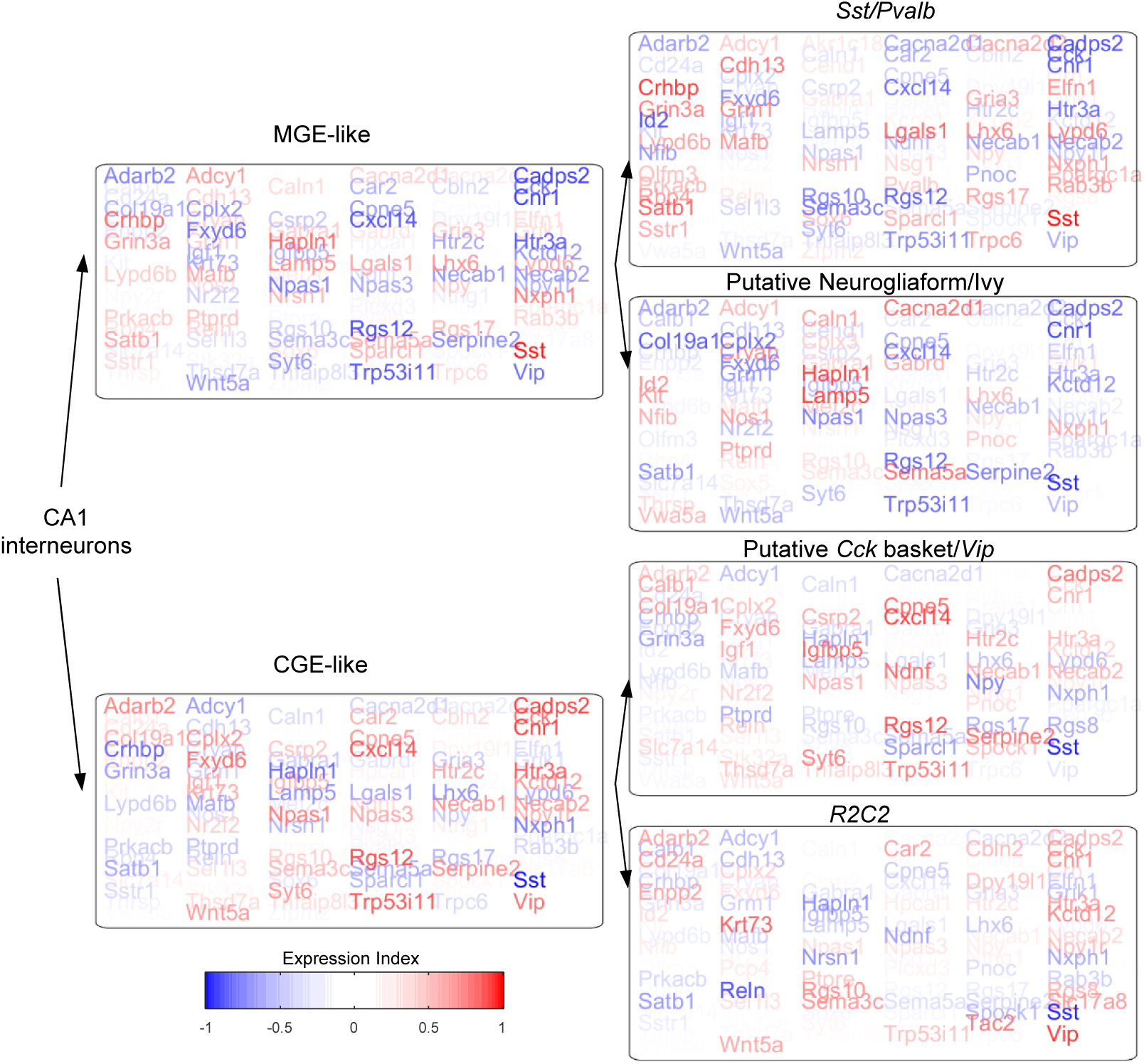
Gene expression in branches at the 1^st^ and 2nd hierarchical levels. Each box represents a classification branch, with the relative expression index for selected genes represented in pseudocolor. Red indicates higher mean expression inside this branch than in the remaining population, blue indicates lower mean expression, and white indicates no difference (see Experimental Procedures for mathematical definition). The genes shown here are a selected set whose expression together prominently differentiates all classes, and are not restricted to members of the gene teams.

**Supplementary Figure 4, associated with Figure 4.**
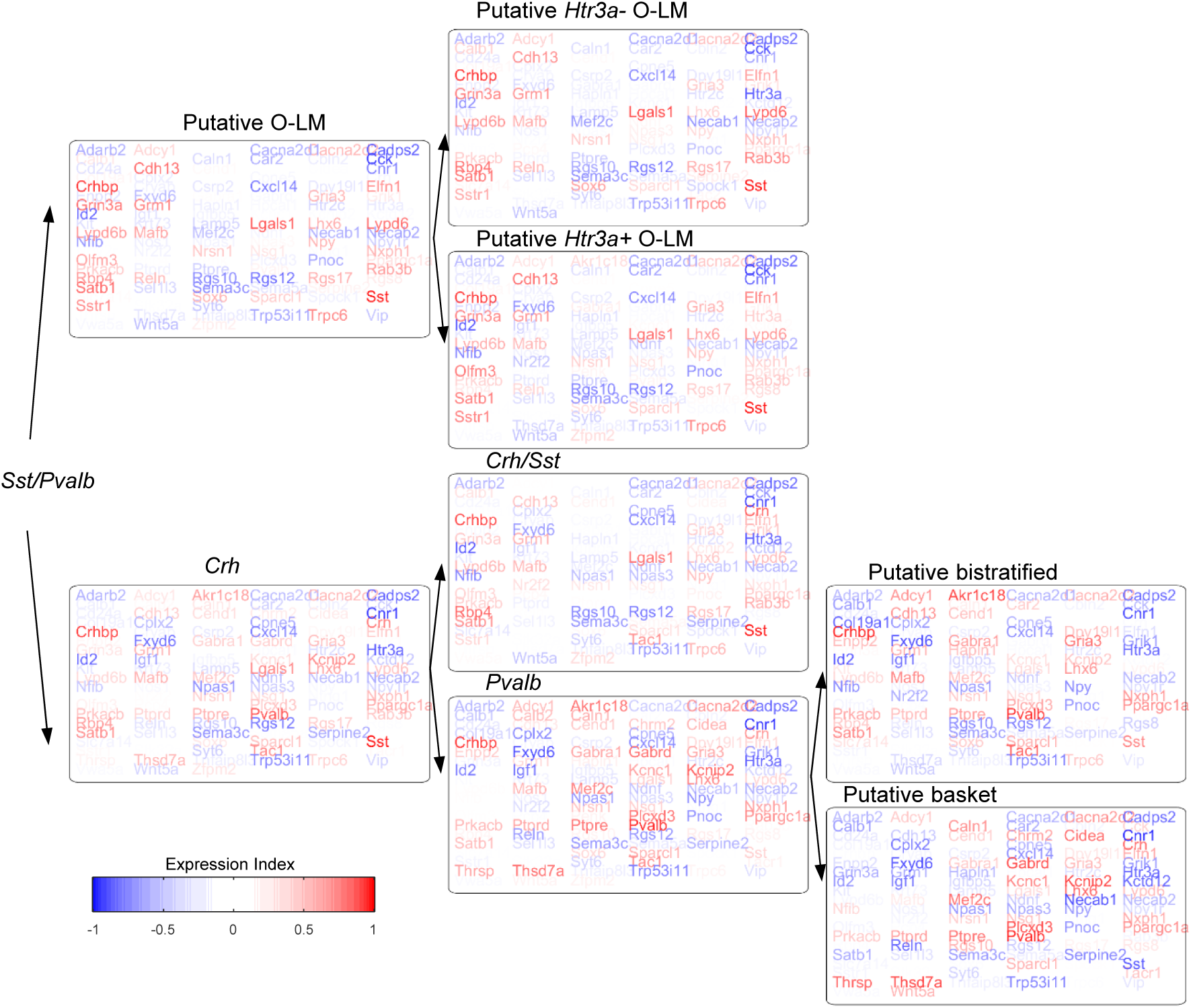
Gene expression in 3rd to 5th level branches of the *Sst/Pvalb* subtree.

**Supplementary Figure 5, associated with Figure 4.**
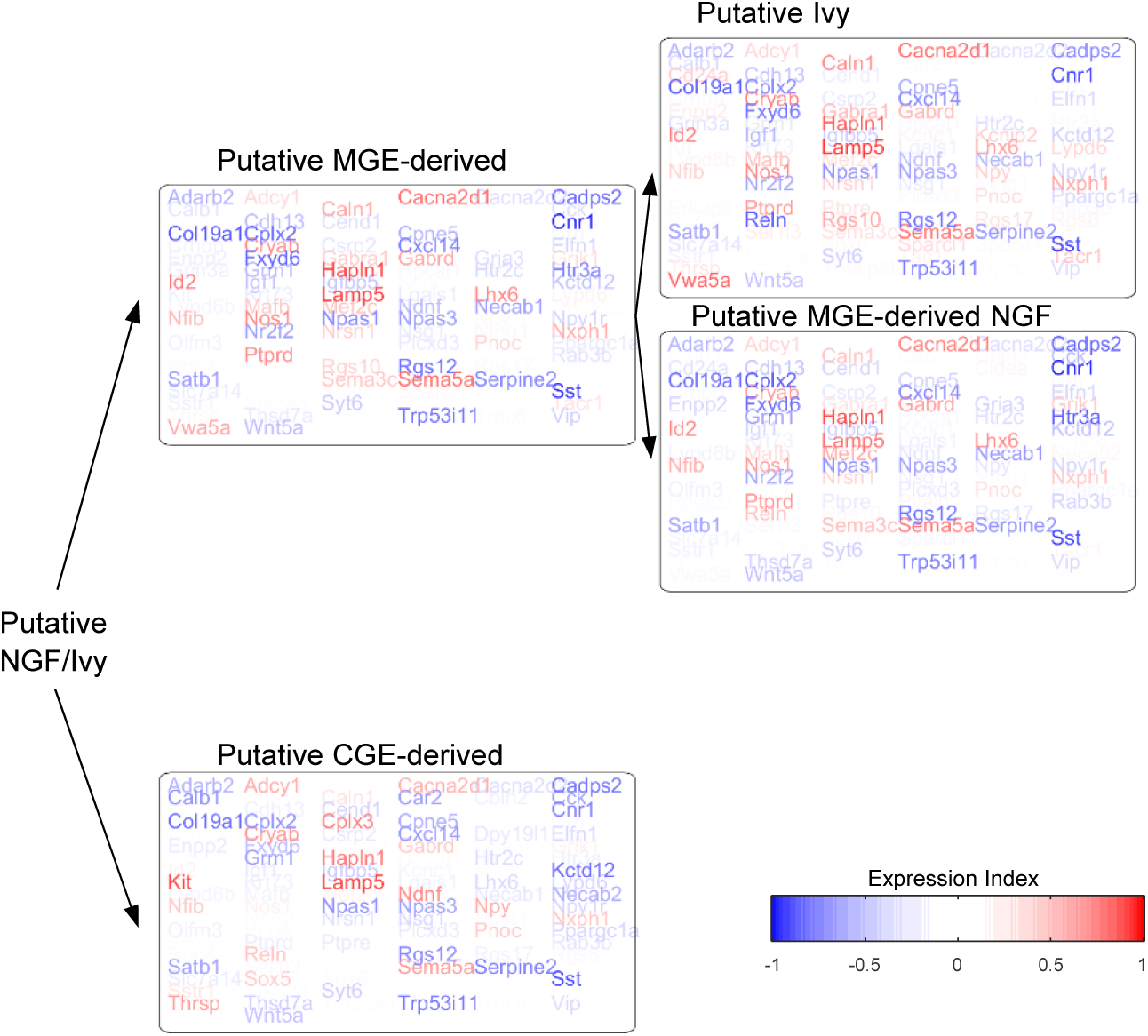
Gene expression in 3^rd^‐4^th^ level branches at of the putative Neurogliaform/Ivy subtree.

**Supplementary Figure 6, associated with Figure 4.**
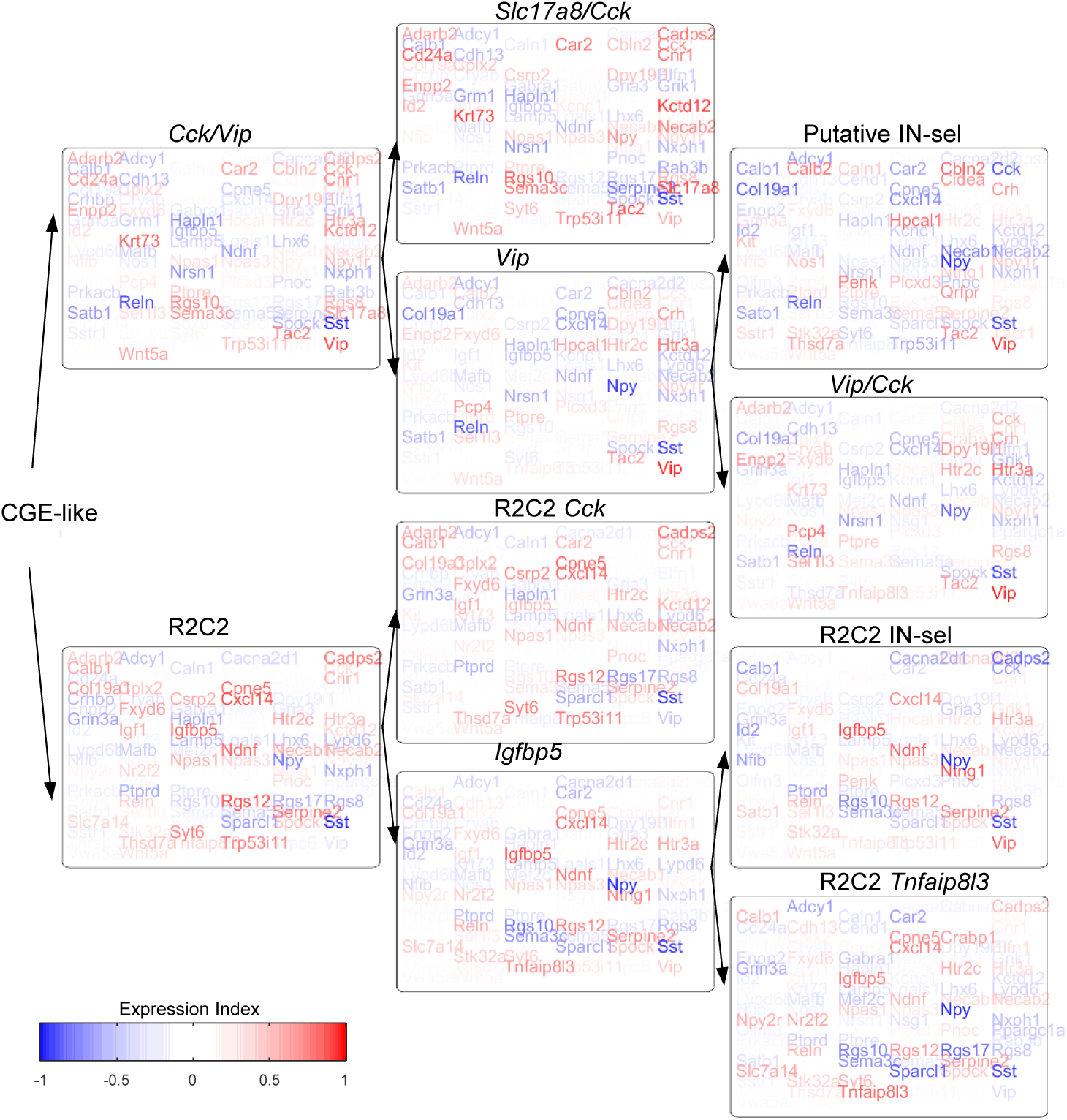
Gene expression in 2rd‐4th level branches of the CGE‐like subtree.

**Supplementary Figure 7.**
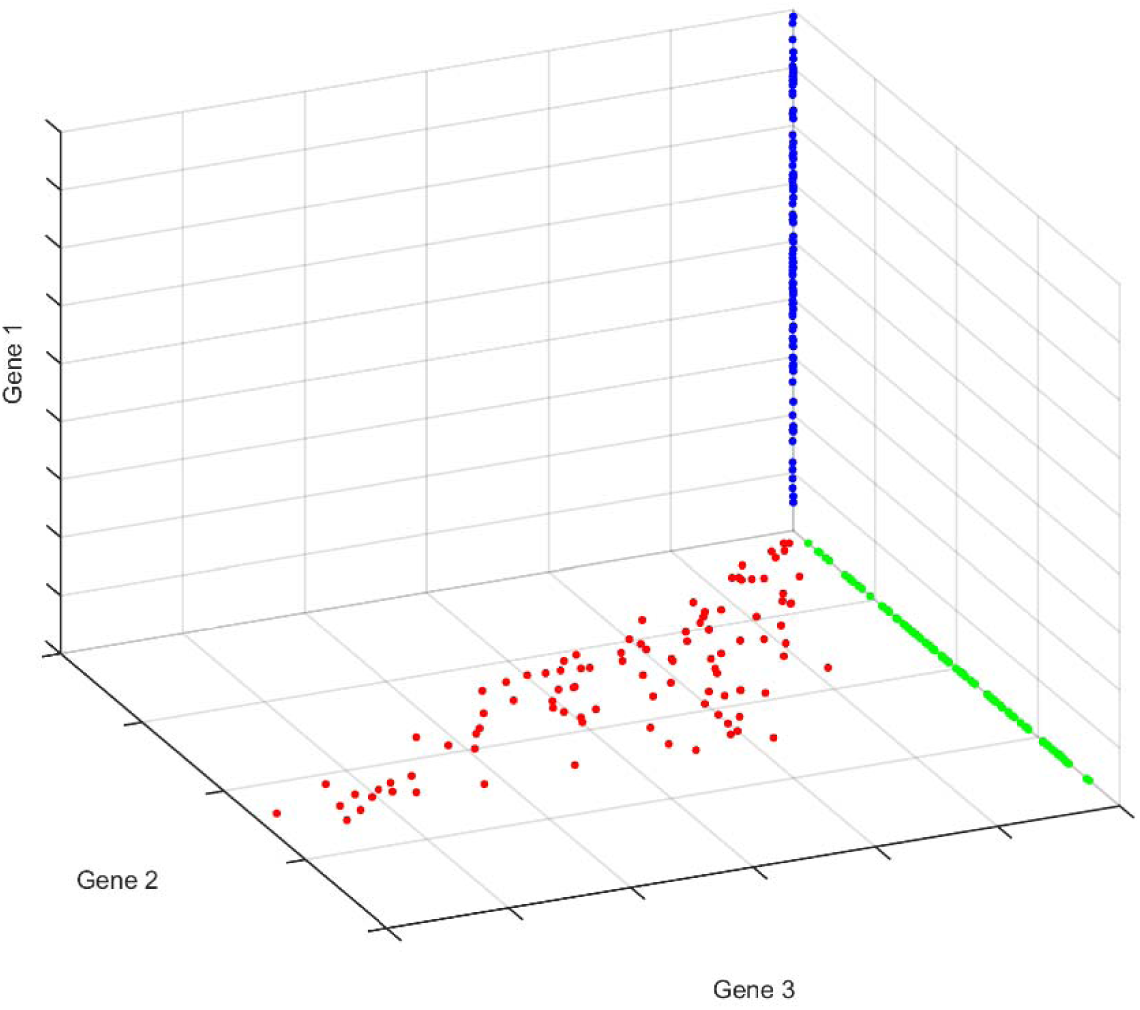
Cartoon illustration of how gene expression varies within and between cell classes. Colored dots represent hypothetical expression patterns of individual cells in a 3-dimensional subspace of expression vectors. The blue class expresses only gene 1, the green and red classes express gene 2 but not gene 1, and the red class additionally expresses gene 3. We hypothesize that absolute gene expression levels are variable along a continuum within each class (modulated for example by neuronal activity, sleep, and plasticity), but that cells will rarely if ever “jump” between classes by absolutely silencing some genes while unsilencing others.

